# Tanycytic annexinA1-containing extracellular vesicles control thermogenesis by orchestrating microglial and neuronal functions

**DOI:** 10.1101/2025.10.12.681867

**Authors:** Rafik Dali, David Lopez-Rodriguez, Irina Kolotuev, Chaitanya Gavini, Tamara Deglise, Emmanuel Di Valentin, Antoine Rohrbach, Jeanne Buchs, Judith Estrada-Meza, Virginie Mansuy Aubert, Fanny Langlet

## Abstract

Obesity, a major global health issue, results from disrupted energy balance driven by chronic hypothalamic inflammation and altered intercellular communication. Among the diverse cells orchestrating this regulation, tanycytes—specialized ependymal cells at the brain–blood interface—have emerged as key modulators, yet the molecular mechanisms by which they influence surrounding cells remain poorly understood.

Here, we identify Annexin A1 (ANXA1) as a tanycyte-derived anti-inflammatory signal whose expression, localization, and secretion are dynamically regulated by nutritional state and altered under high-fat diet. During positive energy balance, ANXA1 is secreted in CD9⁺ extracellular vesicles (EV), remodeling hypothalamic networks by altering microglial morphology, synaptic density, and neuronal activation. These EV-mediated effects extend systemically to regulate brown adipose tissue thermogenesis, glucose homeostasis, and overall energy balance.

Our findings reveal a previously unrecognized tanycyte–microglia–neuron signaling axis and highlight EV-mediated glial communication as a potential therapeutic target in obesity-associated neuroinflammation.

## INTRODUCTION

The rising prevalence of being overweight or obese presents a pressing public health challenge for our societies. These conditions significantly increase the risk of various metabolic comorbidities, thereby reducing quality of life and causing a substantial burden on healthcare systems^1^.

Body weight regulation is typically described as a balance between energy intake and expenditure, influenced by complex interactions between genetic and environmental factors^2,3^. The brain – particularly the hypothalamus – plays a central role in this process, as it contains diverse cell types that express genes associated with body mass index regulation^4,5^ and whose expression is influenced by the metabolic environment^3^. To function effectively, neuronal and non-neuronal cells are organized into complex regulatory networks and communicate through a wide range of signaling molecules to coordinate physiological responses^6–8^. The hypothalamic network sequentially senses hormones, metabolites, and nutrients released by peripheral organs or derived from the diet, integrates this metabolic information, and fine-tunes various physiological responses primarily through the endocrine and autonomic nervous systems^9^. Obesity is widely recognized as a consequence of impaired hypothalamic networks and regulatory systems, leading to significant alterations in feeding behavior, glucose homeostasis, and thermoregulation^5,10,11^. Understanding these mechanisms is therefore essential for the development of effective therapies against this metabolic disorder.

Key components within the metabolic hypothalamic network are tanycytes, defined as elongated ependymal cells with distinctive morphology, strategic location, and multifaceted functions^12–15^. Tanycytes line the floor and walls of the third ventricle (3v) and extend long processes into the parenchyma, enabling interactions with neurons, glial cells, and blood vessels^16–18^. Their heterogeneity and dynamic gene expression profile allow them to regulate blood-brain and blood-cerebrospinal fluid (CSF) exchanges, control neurosecretion into the peripheral circulation, transport and sense peripheral metabolic cues, coordinate neuronal functions, and modulate neural circuits through their neural stem cell properties^3,19,20^. However, our understanding of how tanycytes communicate within the hypothalamic network in physiological and pathological conditions remains incomplete. A few studies have highlighted bidirectional communication between tanycytes and neurons in physiological conditions, primarily mediated by metabolites such as ATP^21,22^, lactate^23^, or glutamate^24^, and/or by secreted proteins and peptides such as chemerin^25^ and diazepam binding inhibitor (DBI)^26–28^. Nevertheless, the underlying molecular mechanisms, the full spectrum of tanycyte-derived signals, their specific cellular targets within the hypothalamic network, and how these processes are altered in pathological conditions such as obesity, remain unknown but crucial to understanding the complex role of tanycytes in energy balance regulation.

In this study, we reveal that tanycytes communicate with neural cells through extracellular vesicles (EVs) containing Annexin A1 (ANXA1), an anti-inflammatory protein identified through proteomic analysis. Specifically, ANXA1 is expressed in the ependymal layer of the mediobasal hypothalamus, dynamically regulated by metabolic state, and altered in diet-induced obesity. Through EV-mediated secretion, ANXA1 influences microglial and neuronal activity, forming a communication network that controls thermoregulation, glucose homeostasis, and energy balance.

## RESULTS

### Tanycytes release anti-inflammatory Annexin A1 through extracellular vesicles

To investigate how tanycytes might communicate with other hypothalamic cells, we first examined their ultrastructure using electron microscopy (EM) (Figure 1A-D)^16^. We notably observed multivesicular cargos within tanycyte processes, positioned near the plasma membrane (Figure 1A-B, Supplemental Media 1). These cargos are characterized by multiple internal vesicles enclosed within a single outer membrane, with luminal vesicles ranging from 120 to 200 nm in diameter, and the entire organelle measuring around 750 nm (Figure 1C-D). Even though these measurements are precise, they might not be absolute since the tissue tends to shrink ∼10% during the EM preparation dehydration and embedding steps. Similar multivesicular cargos were found in primary tanycyte cultures (Figure 1E), in which cells maintain the elongated morphology of tanycyte and the expression of canonical markers such as vimentin (Supplemental Figure 1A-F, Supplemental Media 2). Such observations suggest the presence of unconventional secretory pathways mediated by extracellular vesicles (EVs) both *in vivo* and *in vitro*. To further support this hypothesis, we interrogated our scRNAseq dataset to determine whether tanycytes are enriched in genes related to EV biogenesis, machinery, and secretion. This analysis revealed that the ependyma, particularly tanycytes, co-express a high number of genes linked to EV secretion, including those cataloged in ExoCarta^29^ and Vesiclepedia^30^ (Supplemental Figure 1G-I), suggesting that tanycytes are prone to secrete EVs.

**Figure 1.**
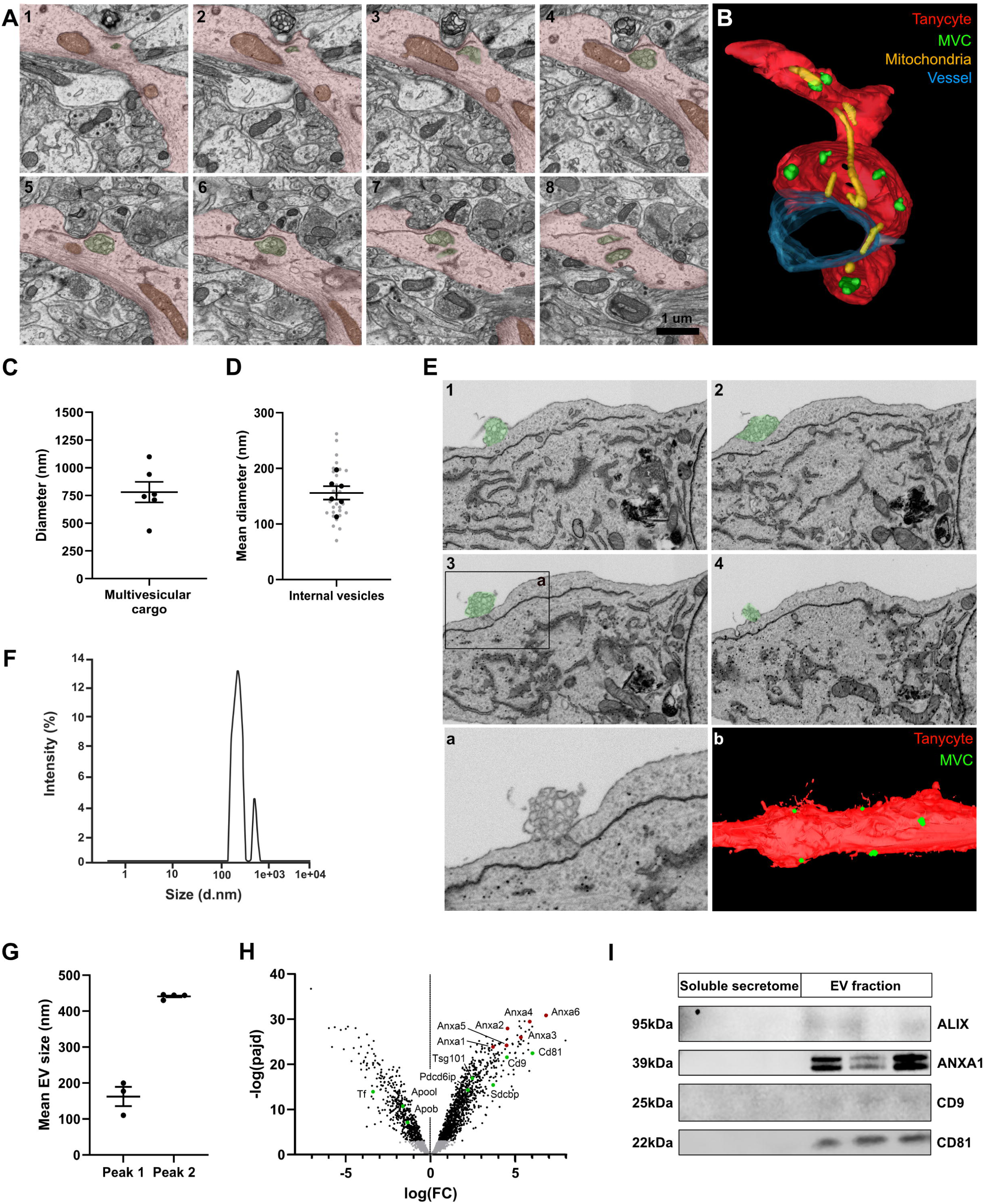
Tanycytes release Annexin A1-containing extracellular vesicles. **(A-B)** Series of scanning electron microscopy micrographs showing the ultrastructure of multivesicular cargos (MVC, green) within tanycyte process (red) (A) and their 3D-representation (B) in the tanycyte process in adult male mice (mitochondria in yellow, blood vessel in blue). **(C-D)** Mean size of multivesicular cargos (C) and internal vesicles (D) observed in tanycyte processes. A total of 6 different MVCs in 2 different tanycytes and 3/6 vesicles per MVC were analyzed. **(E)** Series of scanning electron microscopy micrograph showing the ultrastructure of a multivesicular cargo (green) in primary tanycyte cultures, with a high magnification picture (a, insert in 3) and 3D-representation (b). **(F)** Representative particle size distribution profiles for extracellular vesicles obtained from primary tanycyte culture media by ultracentrifugation analyzed by Nanoparticle Tracking Analysis (NTA). **(G)** Mean size of secreted EV analyzed by NTA. Two technical replicates in 2 different cultures were analyzed (n=3-4). **(H)** Volcano plot displaying enriched proteins in the tanycyte EV *versus* soluble secretome fraction obtained from primary tanycyte culture media. Three technical replicates in 3 cultures were analyzed (n=9 per group). **(I)** Western blot confirming the presence or absence of ANXA1, ALIX, CD9, and CD81 in tanycyte-derived EV fraction and soluble secretome.

EVs carry a diverse array of bioactive molecules, including secreted proteins and peptides, which could play a role in tanycyte-neural cell communication. To next identify the protein composition of tanycyte-derived vesicles, we isolated them from tanycyte culture media using ultracentrifugation^31^. Nanoparticle Tracking Analysis (NTA) revealed a small population of 440 nm vesicles and a larger population of 160 nm vesicles (Figure 1F-G), corresponding to the typical characteristics of large EVs. Proteomic analysis confirmed their identity as EVs, as 92% of the top 100 EV markers —such as CD63, CD9 (already reported in α-tanycytes^16,32^), CD81, and CD82— were present in the tanycyte-derived EV fraction (Supplemental Figure 1J-L, Supplemental Table 1). Additionally, we verified the absence/low amount of non-EV-associated proteins, including those from the TOM complex, CYCS (for mitochondria), RYR, CANX (for the endoplasmic reticulum), and GOLGA2 (for the Golgi apparatus) (Supplemental Table 1). Among the top 50 proteins in tanycyte-derived EVs, the annexin superfamily—comprising 13 members known for their Ca^2+^-binding C-terminal domains that facilitate binding to negatively charged cell membranes^33^ —was prominently represented, including ANXA1, ANXA2, ANXA3, and ANXA6 (Figure 1H, Supplemental Table 1).

By integrating our proteomic data, single-cell RNA sequencing^20,34,35^, and a literature review^36–39^, we selected ANXA1, also known as lipocortin-1^40^, as a key tanycyte candidate for regulating metabolism through EVs. Indeed, within the mediobasal hypothalamus, ANXA1 is selectively expressed in the ependymal region^34,40^, including both α-tanycytes and typical ependymal cells (Supplemental Figure 1M). Recent investigations further underscore ANXA1’s significance in conditions such as obesity and associated comorbidities, including type 2 diabetes^41,42^. Unlike other annexins, ANXA1’s N-terminal interacts with formyl peptide receptors (FPR1 and 2), triggering anti-inflammatory and pro-resolving responses^39^. We finally confirmed by western blot analysis that ANXA1 is specifically secreted in tanycytic EVs: it was detected exclusively in the EV fraction of tanycyte-conditioned medium, alongside EV markers such as CD81, CD9, and ALIX, and was absent from the EV-depleted soluble secretome, confirming the identity of the isolated vesicles (Figure 1I).

### Annexin A1 expression in tanycytes is subgroup-specific and dynamically regulated by the metabolic state

We next mapped ANXA1 expression in the mouse mediobasal hypothalamus using *in situ* hybridization and immunohistochemistry. ANXA1 proteins and mRNAs are localized along the 3v with no detectable signal in the hypothalamic parenchyma (Figure 2A-C). Specifically, ANXA1 protein is detected in tdTomato-expressing ependymal cells and tanycytes (Figure 2A) but not in astrocytes, microglia, blood vessels, and neurons (Supplemental Figure 1N), consistent with previous reports in rat brains^40^. Along the ventrodorsal axis of the 3v, ANXA1 levels are higher in VMH/DMH-tanycytes and ependymal cells compared to ME/ARH tanycytes (Figure 2D). No significant differences were detected along the anteroposterior axis (Figure 2D). Additionally, ANXA1 subcellular localization varies across tanycyte subtypes: in the ARH, ANXA1 is confined to basal endfeet, whereas it is distributed throughout the entire cell in VMH/DMH tanycytes (Figure 2E). Notably, ANXA1 is frequently observed in tanycyte protrusions (Figure 2C) -previously described as potential communication interfaces contacting neurons, glial cells, and blood vessels^16^.

**Figure 2.**
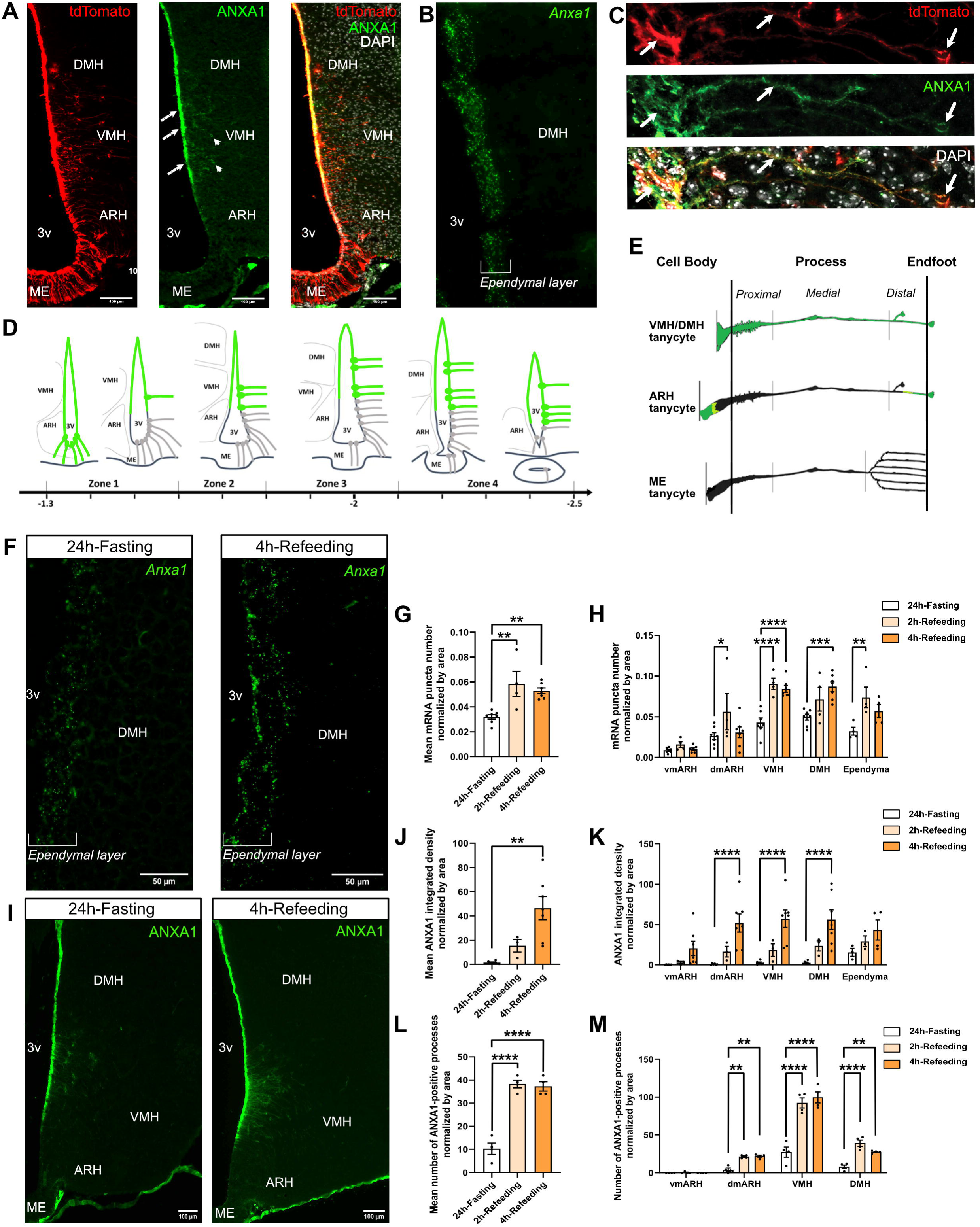
Tanycyte ANXA1 expression is modulated by the metabolic state. **(A)** Representative images of ANXA1 immunostaining (green) along the ventricle (arrows). ANXA1 staining (green) colocalizes with tdTomato-positive tanycytes (red). Cell nuclei are counterstained using DAPI (white). **(B)** Representative images of *Anxa1* mRNA expression (green) along the third ventricle (brackets). **(C)** Representative images of ANXA1 immunostaining (green) along tdTomato-positive tanycyte process (red) along the VMH. **(D)** Schematic representation of ANXA1 expression (green) along the third ventricle on the ventrodorsal and anteroposterior axis based on *in situ* hybridization and immunohistochemistry analysis. **(E)** Schematic representation of ANXA1 subcellular localization within each tanycyte subtype based on immunohistochemistry analysis. **(F-H)** Representative images of *Anxa1* mRNA expression (green, F) along the third ventricle during fasting-refeeding paradigm and quantification on the entire region (G) and the anteroposterior axis (H) (n=4-7 mice/group in two cohorts). **(I-K)** Representative images of ANXA1 expression (green, I) along the third ventricle during fasting-refeeding paradigm and quantification on the entire region (J) and the anteroposterior axis (K) (n=3-7 mice/group in two cohorts) **(L-M)** Quantification of ANXA1 distribution in tanycyte processes during fasting-refeeding paradigm on the entire region (L) and the anteroposterior axis (M) (n=4 mice/group in one cohort) during fasting-refeeding paradigm. ARH, arcuate nucleus; DMH, dorsomedial nucleus; VMH, ventromedial nucleus; 3v, third ventricle. *p < 0.05, \*\**p* < 0.01, \*\*\**p* < 0.001, \*\*\*\**p* < 0.0001.

To determine whether ANXA1 is regulated by the metabolic state, we examined its expression under physiological (fasting-refeeding paradigm) (Figure 2F-M) and pathological (high-fat diets) conditions (Supplemental Figures 2 and 3). Refeeding after a 24h-fasting induced a robust increase in ANXA1 expression (Figure 2F-K) and its redistribution along tanycytes processes (Figure 2L-M), particularly in the dmARH, VMH, and DMH. High-fat diets are known to trigger hypothalamic inflammation, a key driver of obesity and metabolic dysfunctions characterized by microglial and astrocyte activation and the release of pro-inflammatory cytokines, such as IL-1β, TNF-α, and IL-6^43^. Given the anti-inflammatory and pro-resolving functions of ANXA1^39^, we hypothesized that its expression may be dynamically regulated across the distinct phases of diet-induced neuroinflammation (Supplemental Figures 2 and 3, respectively). While short-term (a few hours) exposure to a high-fat/high-sucrose (HFHS) diet does not induce changes in microglial and astroglial density and morphology, ependymal ANXA1 expression increases after 3h (Supplemental Figure 2B-F). Medium-term exposure (several days) to HFHS diet induces a marked astrocytic activation on day 3, followed by a decrease on day 14 and a rebound on day 28, as previously described^44^, without any changes in microglial activation or ANXA1 expression along the ventricular wall (Supplemental Figure 2G-K). Finally, long-term (8 weeks) exposure to HFHS diet induces an increase in astroglial density and morphology associated with a decrease in ANXA1 expression along the ventricle (Supplemental Figure 2L-S). Interestingly, a 60% high-fat (60HF) diet induces a more pronounced effect on microglia and ANXA1 expression (Supplemental Figure 3). Particularly, medium-term exposure to 60HF diet induces a marked microglial activation on day 3, followed by a return to baseline on day 14^44^, with ANXA1 expression levels following the same pattern (Supplemental Figure 3G-K). After 8 weeks of 60HF diet, ANXA1 expression decreases, associated with microglial and astrocytic activation (Supplemental Figure 3L-T). Such a decrease in tanycyte ANXA1 expression after long-term HFD is further validated by TRAP-sequencing done in tanycytes from normal chow *versus* 12 weeks HFD mice^45^.

**Figure 3.**
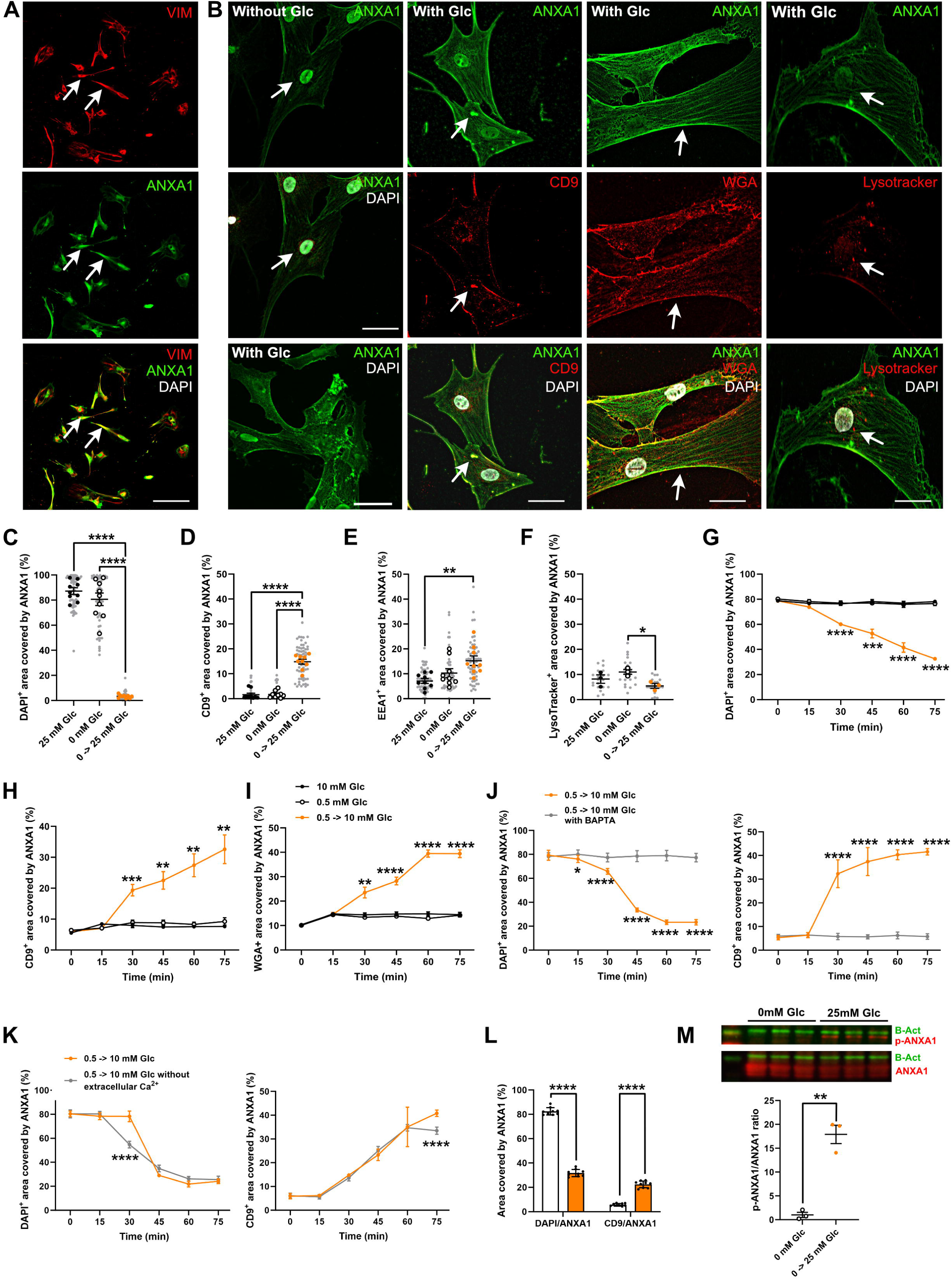
ANXA1 relocalizes within the cell upon glucose treatment. **(A)** Representative images of ANXA1 immunostaining (ANXA1 in green, Vimentin in red) in primary tanycyte cultures in the presence of glucose. **(B)** Representative images of ANXA1 (green) within nuclei (white, DAPI-positive), multivesicular bodies (red, CD9-positive), cell membrane (red, WGA-positive), and lysosomes (red, Lysotracker-positive) in primary tanycyte cultures following glucose treatment. **(C-F)** Quantifications of ANXA1 within nuclei (DAPI-positive, C), multivesicular bodies (CD9-positive, D), early endosomes (EEA1-positive, E), and lysosomes (Lysotracker-positive, F) in primary tanycyte cultures following glucose treatment for 1h after glucose deprivation for 24h. 6-7 tanycytes per well were analyzed in three technical replicates in three cultures (n=9 per condition; light gray points indicate each tanycyte analyzed). **(G-I)** Time-course quantification of ANXA1 redistribution from the nucleus (G) to CD9-positive vesicles (H) and cell membrane (I) following glucose exposure (after glucose deprivation for 24h). 6-7 tanycytes per well were analyzed in three technical replicates in three cultures (n=9 per condition). **(J-K)** Time-course quantification of ANXA1 redistribution from the nucleus to CD9-positive vesicles following glucose exposure (after glucose deprivation for 24h) in the absence or presence of BAPTA (J) and free-calcium Krebs medium (K). 6-7 tanycytes per well were analyzed in three technical replicates in three cultures (n=9 per condition). **(L)** ANXA1 redistribution from the nucleus to CD9-positive vesicles in the absence or presence of thapsigargin during glucose deprivation in free-calcium Krebs medium. 6-7 tanycytes per well were analyzed in three technical replicates in three cultures (n=9 per condition). **(M)** ANXA1 Tyr21 phosphorylation in primary tanycyte cultures following glucose treatment for 2h after glucose deprivation for 24h. 3 wells were analyzed in one culture (n=3 per condition). *p < 0.05, \*\**p* < 0.01, \*\*\**p* < 0.001, \*\*\*\**p* < 0.0001.

### Insulin and glucose control Annexin A1 expression and localization in tanycytes

To investigate how the metabolic conditions regulate ANXA1 expression and localization within tanycytes, we used primary tanycyte cultures and exposed them to various hormones and nutrients, mimicking the distinct energy states. ANXA1 was detected in nearly all tanycytes, with strong nuclear distribution under basal conditions (Figure 3A). While glucose, dexamethasone, or insulin alone does not significantly impact *Anxa1* mRNA expression (Supplemental Figure 4A-C), treatment with cAMP/dexamethasone (simulating the fasting state) reduces protein levels at 2h and *Anxa1* transcript at 4h (Supplemental Figure 4D-E). The addition of insulin to this treatment (mimicking the early refeeding state) results in a 1.5-fold increase in ANXA1 expression after 2 hours (Supplemental Figure 4D-E).

Although glucose does not modulate *Anxa1* mRNA expression (Supplemental Figure 4A), it induces striking changes in ANXA1 protein localization within tanycytes (Figure 3B) and a slight decrease in ANXA1 protein cellular content after 4h (Supplemental Figure 4F). Under glucose-deprived conditions *in vitro*, ANXA1 remains predominantly nuclear, whereas glucose treatment leads to its redistribution to early EAA1-positive endosomes, CD9-positive multivesicular cargos, and plasma membrane, avoiding lysosomal compartments (Figure 3B-F) —similar to what has been seen in other cell types (available on Gene Cards: ANXA1). Notably, this redistribution is evident from 15mM glucose (Supplemental Figure 4G-I) and within 30 minutes (Figure 3G-I), coinciding with the appearance of CD9-positive vesicles (Supplemental Figure 4J), highlighting a dose- and time-dependent effect.

Given that ANXA1 binds membrane phospholipids in the presence of Ca²⁺ ^39^ and glucose activates Ca²⁺ signaling in tanycytes^21,22^, we next explored the role of Ca^2+^ in ANXA1 redistribution. Intracellular calcium chelation with BAPTA effectively prevents ANXA1 redistribution following glucose treatment (Figure 3J). In contrast, glucose-induced ANXA1 relocalization is unaffected by incubation in Ca²⁺-free Krebs medium, highlighting the critical role for intracellular, rather than extracellular calcium (Figure 3K). Artificially increasing intracellular calcium with thapsigargin is also sufficient to trigger ANXA1 redistribution within 15 minutes (Figure 3L). Finally, we examined ANXA1 phosphorylation at tyrosine 21 in its N-terminal domain^46^ - a post-translational modification known to facilitate membrane docking and EV packaging^47^. Glucose treatment induces phosphorylation at this site in tanycytes (Figure 3M), supporting a model in which glucose mobilizes ANXA1 through intracellular calcium signaling and post-translational modification, enabling its incorporation into multivesicular compartments.

### Tanycytic Annexin A1 modulates energy balance and thermoregulation

To examine the physiological relevance of tanycytic ANXA1, we generated tanycyte-specific *Anxa1* knockout (TanAnxa1^−/−^ mice) mice by infusing AAV1/2-*Tshr*-Cre into the dorsal third ventricle of adult *Anxa1*^F/F^. This model eliminates full-length ANXA1, including both N- and C-terminal domains, and mirrors the downregulation observed under chronic HFD exposure (Supplemental Figures 2 and 3). Deletion efficiency was confirmed along the third ventricle by quantifying mRNA levels (Supplemental Figure 5A-B). Under basal conditions, TanAnxa1^−/−^ mice show no differences in body weight or glycemia (Supplemental Table 2), but exhibit a significant increase in core body temperature starting four weeks post-deletion (Figure 4A). After 24h-fasting, these mice showed a slight reduction in food intake and core body temperature during refeeding (Figure 4B-D). In metabolic cages, these mice display reduced energy expenditure per hour and reduced cumulative food intake (Figure 4E-F), resulting in no change in energy balance (Figure 4G). Notably, the energy expenditure defect is abolished under thermoneutral conditions (Supplemental Figure 5G), suggesting altered sympathetic tone. Thermoregulation and counterregulation responses to hypoglycemia were further impaired in TanAnxa1^−/−^ mice. Upon 2-deoxyglucose (2DG) injection, body temperature remains unchanged, but glycemic excursions are blunted (Figure 4H-I). Following norepinephrine administration, control mice show a typical drop in temperature – likely due to vasodilatation – whereas knockout mice do not (Figure 4J-K). Once again, glucose excursions are diminished in TanAnxa1^-/-^ mice (Figure 4J-K). During cold exposure, TanAnxa1^−/−^ mice exhibit a higher glycemia and a rapid drop in body temperature (Figure 4L-M), consistent with defective thermogenic compensation. Altogether, these data indicate an altered thermoregulation in TanAnxa1^-/-^ mice likely associated with BAT hyperactivation. Indeed, BAT histological analysis revealed increased thermogenic activity in TanAnxa1^−/−^ mice, characterized by numerous small lipid droplets and upregulation of sympathetic nervous system-responsive genes such as *Dio2* (Supplemental 5I-K). To confirm BAT hyperactivation, we used a β3-adrenergic receptor antagonist to inhibit BAT activity: this treatment induced a marked drop in body temperature in TanAnxa1^⁻/⁻^ mice, without affecting glycemia (Figure 4N-O).

**Figure 4.**
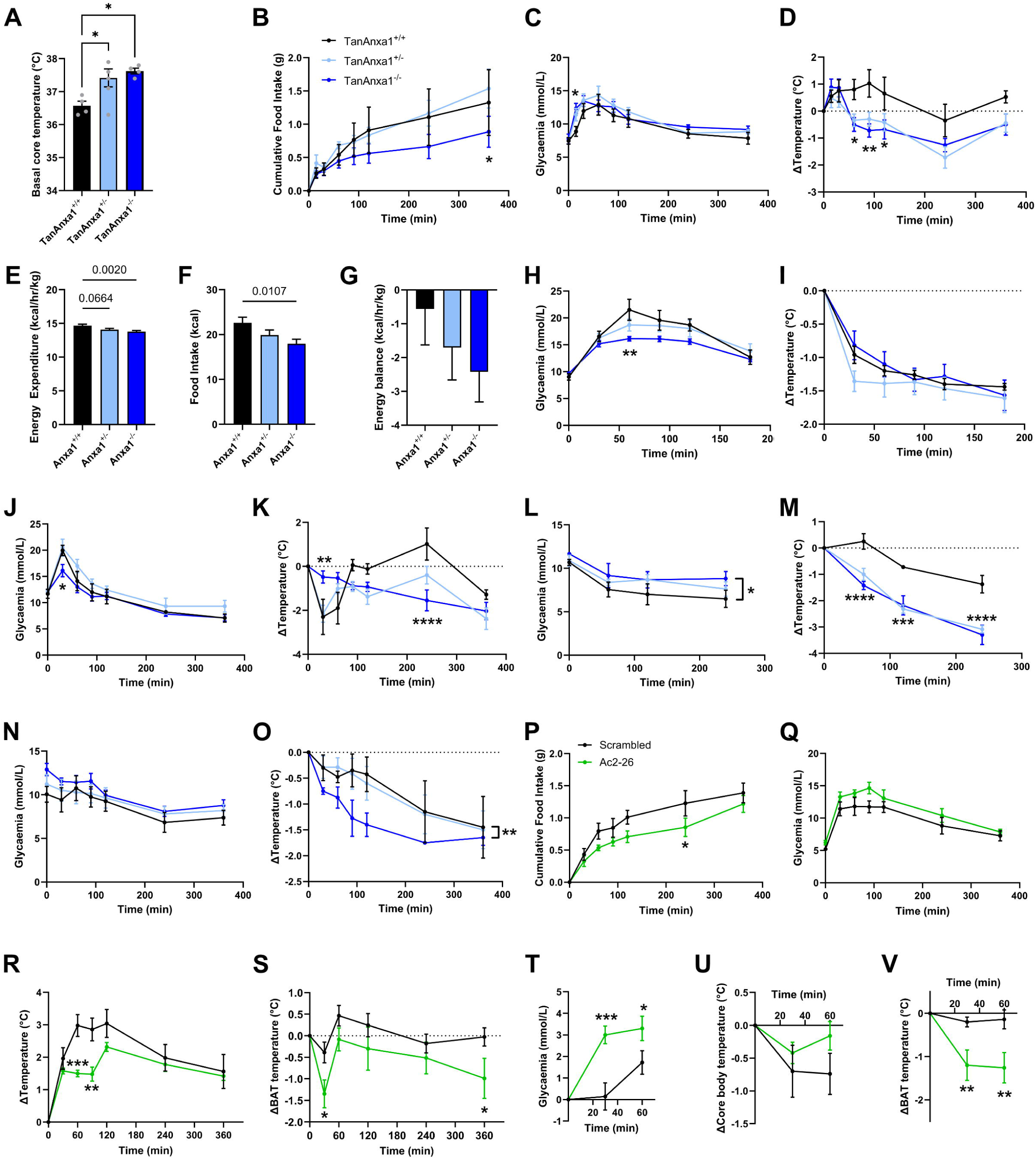
Tanycyte ANXA1 impacts energy balance, glucose homeostasis, and thermoregulation. **(A)** Basal core temperature in CTL versus TanAnxa1^−/−^ mice. **(B-D)** Cumulative food intake (B), glycemia (C), and delta core temperature (D) in CTL *versus* TanAnxa1^−/−^ mice during the refeeding. **(E-G)** Energy expenditure per hour (E), cumulative food intake (F), and energy balance per hour (G) in metabolic cages in CTL *versus* TanAnxa1^−/−^ mice. **(H-I)** Glycemia (H) and delta core temperature (I) following 2DG injection in CTL *versus* TanAnxa1^−/−^ mice. **(J-K)** Glycemia (J) and delta core temperature (K) following norepinephrine injection in CTL *versus* TanAnxa1^−/−^ mice. **(L-M)** Glycemia (L) and delta core temperature (M) during cold exposure in CTL *versus* TanAnxa1^−/−^ mice. **(N-O)** Glycemia (N) and delta core temperature (O) following β3 adrenergic antagonist in CTL *versus* TanAnxa1^−/−^ mice. **(P-S)** Cumulative food intake (P), glycemia (Q), delta core temperature (R), and delta BAT temperature (S) in 24h-fasted mice injected with Ac2-26 active peptide just before the refeeding. **(T-V)** Glycemia (T), delta core temperature (U), and delta BAT temperature (V) in fed mice injected with Ac2-26 active peptide. *p < 0.05, \*\**p* < 0.01, \*\*\**p* < 0.001, \*\*\*\**p* < 0.0001.

To further validate the functional role of ANXA1 signaling, we next infused the N-terminal ANXA1 analog Ac2-26 intracerebroventricularly (i.c.v.) in adult mice under fasted or fed conditions. In fasted mice, Ac2-26 reduces food intake, measured manually (Figure 4P) or using metabolic cages (Supplemental Figure 5L and N), without affecting glycemia (Figure 4Q) or energy expenditure (Supplemental Figure 5M and O). Additionally, Ac2-26 causes a reduction in core body temperature during refeeding, which correlated with a significant decrease in brown adipose tissue (BAT) temperature (Figure 4R-S). In fed mice, Ac2-26 similarly reduces BAT temperature 1h after injection without affecting core body temperature and increases glycemia (Figure 4T-V). Once again, BAT activity seems to be affected. Histological analysis showed fewer but larger lipid droplets, and lipogenic gene expression (i.e., *Srebf1, Fasn*) was upregulated (Supplemental Figure 5P-R), consistent with reduced BAT thermogenesis.

Altogether, these data show that tanycytic ANXA1 acts as a late post-prandial signal that maintains glycemia and reduces temperature.

### Tanycytic ANXA1 engages microglia and neurons via FPR signaling

To investigate the downstream targets of tanycyte-derived ANXA1, we examined the expression of its known receptors, FPR1 and FPR2^39^, in the mouse hypothalamus. The mouse HypoMap dataset^34^ revealed that *Fpr1* and *Fpr2* are mainly expressed by microglia and VMH neurons, respectively (Supplemental Figure 6A-B). Within the neuronal population, *Fpr2* is expressed in several neuropeptidergic subtypes involved in energy balance regulation, including SF1, POMC, and NPY neurons (Supplemental Figure 6C). Using *in situ* hybridization and immunohistochemistry, we confirmed that FPR1 is broadly expressed throughout the mediobasal hypothalamus, with significantly higher levels in nuclei adjacent to the third ventricle—namely the arcuate (ARH), ventromedial (VMH), and dorsomedial (DMH) hypothalamic nuclei—compared to the lateral hypothalamus (LH) (Supplemental Figure 6D, F, H). In contrast, FPR2 exhibited a more diffuse expression pattern, detectable in both the tanycyte layer and the surrounding parenchyma, albeit at lower levels than FPR1 and without significant regional differences across hypothalamic nuclei (Supplemental Figure 6E, G, I). Expression levels of both FPR1 and FPR2 remained unchanged upon fasting–refeeding paradigm (Supplemental Figure 6J-N).

To test the functional effects of ANXA1 signaling on microglia and neurons, we infused the Ac2-26 peptide (an active N-terminal peptide of ANXA1 binding FPRs) into the lateral ventricle of fasted mice. Two hours post-infusion, c-Fos and pSTAT3 (a signaling pathway induced by Ac2-26^48^) were elevated in the dmVMH and DMH (Figure 5A-F), regions involved in glucose counterregulatory response^49^ and thermogenic control^50^, respectively. This activation was mostly restricted to HuC/D^+^ neurons (Figure 5C, F), notably in SF1^+^ neurons, while POMC neurons remained unaffected (Supplemental Figure 7A-E). Additionally, glutamatergic terminal density increased in the hypothalamic parenchyma following Ac2-26 treatment (Figure 5G-H), suggesting synaptic plasticity. Strikingly, Ac2-26 also induced morphological changes of microglia, increasing their number and ramification in ARH, VMH, and DMH (Figure 5I-K). This effect on microglial function was further confirmed in primary microglia cultures: Ac2-26 treatment reduced CD68 content and decreased synaptosome uptake (Supplemental Figure 7F-I), indicating a shift toward a less phagocytic phenotype. To further determine whether tanycytic ANXA1 may induce such an effect on microglia, we analyzed TanAnxa1^−/−^ mice under fed conditions. Strikingly, *Anxa1* depletion in tanycytes also induced morphological changes of microglia, slightly increasing their number while decreasing their ramification in ARH, VMH, and DMH (Figure 5L-N), indicating a shift toward a pro-inflammatory phenotype^51^. These findings suggest that tanycyte-derived ANXA1 modulates hypothalamic thermoregulation and energy balance by modulating microglia and neuronal functions.

**Figure 5.**
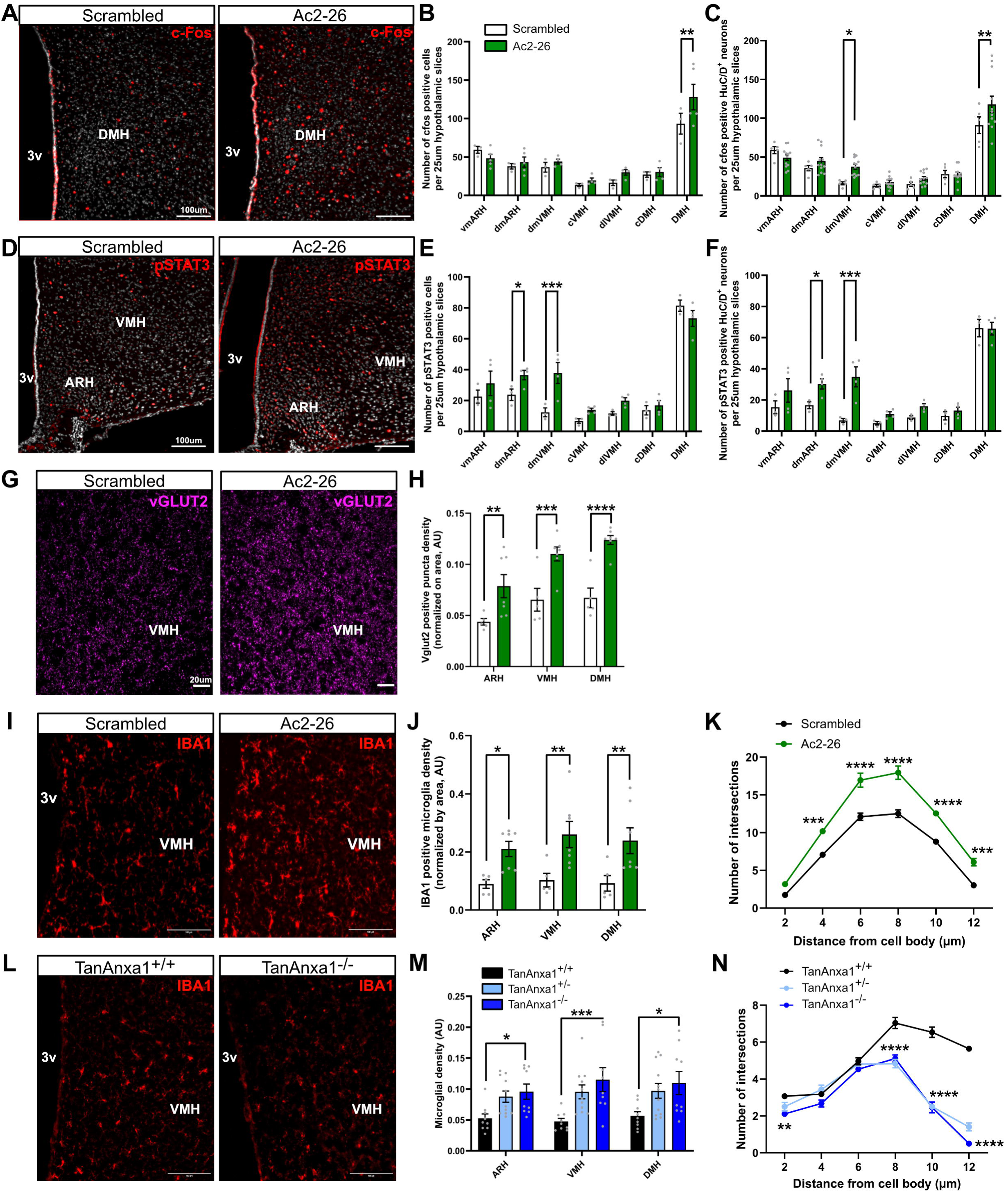
ANXA1 analog Ac2-26 peptide modulates neuronal and microglial functions in the mediobasal hypothalamus. **(A-C)** Representative images (A) and quantifications of c-Fos activation (B-C) in the mediobasal hypothalamus following Ac2-26 i.c.v. infusion in fasted mice. The quantifications indicate the total amount of activated cells (B) and the amount of activated HuC/D-positive neurons (C) per 25 μm hypothalamic slice in two different cohorts. **(D-F)** Representative images (D) and quantifications of pSTAT3 activation (E-F) in the mediobasal hypothalamus following Ac2-26 i.c.v. infusion in fasted mice. The quantifications indicate the total amount of activated cells (E) and the amount of activated HuC/D-positive neurons (F) per 25 μm hypothalamic slice in one cohort. **(G-H)** Representative images (G) and quantifications (H) of VGLUT2-positive synaptic terminals following Ac2-26 i.c.v. infusion in fasted mice. **(I-K)** Representative images (I) and quantifications of microglial density (J) and ramifications (K) following Ac2-26 i.c.v. infusion in fasted mice. **(L-N)** Representative images (L) and quantifications of microglial density (M) and ramifications (N) following Ac2-26 i.c.v. infusion in TanAnxa1^-/-^ mice. *p < 0.05, \*\**p* < 0.01, \*\*\**p* < 0.001, \*\*\*\**p* < 0.0001.

### Microglia depletion rescues thermoregulatory defects in TanAnxa1*^−/−^* mice

To finally determine whether microglia activation mediates the increase in core temperature and the alteration in thermoregulation in TanAnxa1^−/−^ mice, we depleted their microglia using PLX5622, a CSF1R inhibitor (Figure 6). As previously observed, loss of tanycytic ANXA1 initially led to increased basal core temperature, progressive weight gain, and impaired thermogenic responses during acute cold exposure (Figure 6A-C). Remarkably, microglia ablation completely normalized basal temperature and restored cold-induced thermogenesis in these mice (Figure 6D-F), indicating that microglia are critical downstream effectors of tanycytic ANXA1 signaling. Our findings reveal that tanycyte-derived ANXA1 modulates thermoregulatory circuits via microglial reprogramming, revealing an important role for interglial communication in hypothalamic metabolic control.

**Figure 6.**
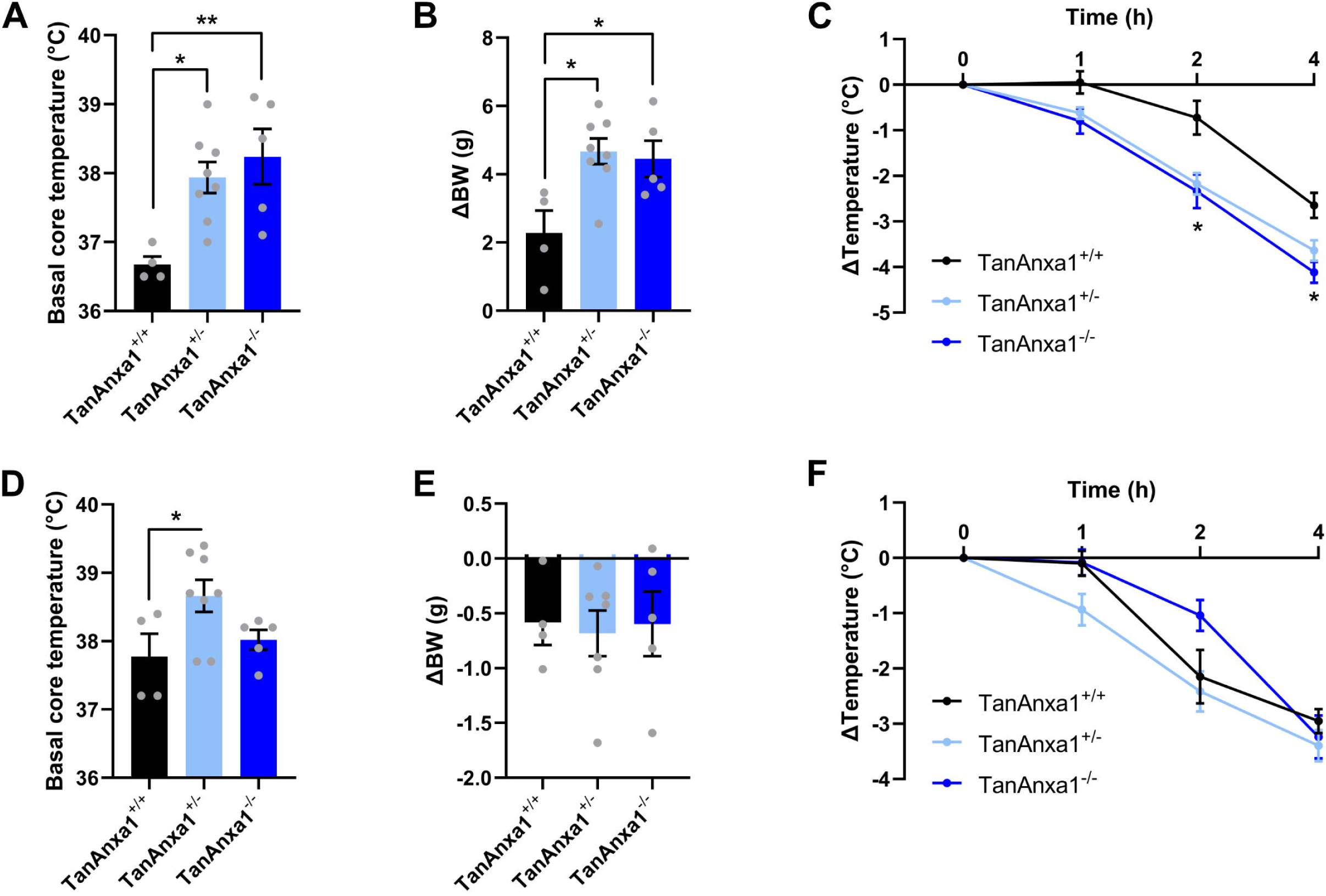
Microglial depletion corrects temperature regulation in TanAnxa1^-/-^ mice. **(A-C)** Basal core temperature (A), delta body weight (B), and delta core temperature in response to cold exposure (C) in CTL *versus* TanAnxa1^−/−^ mice 4 weeks after the deletion. **(D-F)** Basal core temperature (D), delta body weight (E), and delta core temperature in response to cold exposure (F) in CTL *versus* TanAnxa1^−/−^ mice 2 weeks after microglia depletion (in the same cohort as in panel A-C). *p < 0.05, \*\**p* < 0.01, \*\*\**p* < 0.001, \*\*\*\**p* < 0.0001.

## DISCUSSION

This study identifies EVs as a previously unrecognized mode for tanycyte-to-microglia communication in the hypothalamus. These EVs carry the anti-inflammatory protein ANXA1, whose expression and secretion are modulated by metabolic state. Under positive energy balance, ANXA1 redistributes along tanycytic processes, whereas chronic high-fat diet leads to its downregulation. Functionally, tanycyte-derived ANXA1 influences microglial phenotype and neuronal activity in key hypothalamic nuclei, ultimately modulating energy balance, glucose homeostasis, and thermoregulation.

Tanycytes are increasingly recognized as essential components within hypothalamic networks, contributing to the regulation of neuroendocrine functions and systemic metabolism^8,13^. Their role is largely supported by their unique processes, which extend deep into the hypothalamic parenchyma and contact various cell types, including neurons, blood vessels, and glial cells^16–18^. However, despite this neuroanatomical clue, tanycyte-mediated intercellular communication within the hypothalamus remains poorly understood. To date, most studies have focused on interactions with arcuate neurons^52^ via metabolite signaling, such as lactate^23^ and ATP^21,22^. Here, our findings expand this paradigm by demonstrating a tripartite tanycyte-microglia-neuron communication through EVs. This finding not only highlights the complexity of hypothalamic cellular networks but also reveals EV biogenesis and secretion as part of the molecular toolkit of tanycytes. While already suggested through the visualization of multivesicular bodies by electron microscopy^14,15^ and CD9 expression^32,53^, the identification of EVs as carriers of tanycyte-derived signals opens new perspectives for understanding and modulating hypothalamic intercellular communication. Indeed, EVs carry diverse bioactive molecules, including proteins, nucleic acids, and lipids, which influence a wide range of physiological and pathological processes in recipient cells^54^. Under physiological conditions, EV-mediated signaling between neurons and glial cells regulates neuronal activity, modulates immune responses, and helps maintain brain homeostasis^54^. In the context of neuroinflammation, EVs are key messengers among immune cells, including microglia, by diffusing proinflammatory cytokines and chemokines, thereby propagating inflammatory signals throughout the brain^54^. Thus, this dynamic and multifunctional interplay through EVs highlights a highly integrated network, in which each cell type contributes to the functional synergy, forcing us to refine our methodologies to unravel the complexities of hypothalamic regulation of energy balance.

Among the diverse EVs carrying bioactive molecules, we further demonstrate that tanycyte-derived EVs contain substantial amounts of ANXA1. This calcium-dependent phospholipid-binding protein is known to participate in various membrane-related processes, including the EV aggregation/biogenesis^55^, by interacting with phospholipids and cytoskeletal components^39^. While previous studies have reported ANXA1 expression in neurons, astrocytes, microglia, and endothelial cells of the blood-brain barrier under stressful conditions such as Alzheimer’s disease lesions^56^, cancer^57^, or ischemia^58,59^, we detected ANXA1 exclusively along the third ventricle in the hypothalamus, even after an 8-week high-fat diet known to induce neuroinflammation^10,44^. This spatial restriction suggests distinct regulatory mechanisms of ANXA1 in metabolism-related processes, likely influenced by metabolic cues such as insulin and glucose. Indeed, we showed that insulin increases *Anxa1* expression in tanycytes, while glucose redistributes ANXA1 along tanycyte processes via a calcium-dependent and post-translational regulation. Similar metabolic regulations have been described in the literature, particularly in pancreatic β-cells, where ANXA1 is co-secreted with insulin in response to glucose. Indeed, high glucose concentrations enhance ANXA1 phosphorylation in isolated islets, leading to its relocalization into insulin-containing vesicles and its secretion^60^. ANXA1 also enhances insulin secretion by modulating the cytoskeleton and PKC activity in pancreatic β-cells^61^. In the hypothalamus, once secreted by tanycytes, ANXA1 impacts neuronal and glial functions in the neighboring parenchyma, mainly in the DMH and VMH. Notably, the ANXA1-derived peptide Ac2-26 enhances microglial activation and migration, as previously reported^62^. Indeed, its administration promotes a shift in microglial activation and macrophage polarization from a proinflammatory (M1) to an anti-inflammatory (M2) state via the formyl peptide receptor^58^ and downregulates pro-inflammatory cytokines through modulation of the NF-κB pathway^63^. Conversely, FPR1 KO reduces microglial activation^64^.

However, it is also worth noting that EVs, whose formation and secretion may be directly facilitated by ANXA1^55^, also contain numerous other bioactive molecules that could also affect parenchymal cells. Furthermore, given their strategic position along the third ventricle, tanycytes could also secrete EVs directly into the CSF, potentially allowing long-range signaling. ANXA1-containing EVs were notably already found in the CSF^65^. In this context, understanding the mechanisms and stimuli that regulate tanycyte EV production, composition, and secretion will provide invaluable insights to decipher their role in hypothalamic function and metabolic control, and to uncover novel therapeutic strategies targeting metabolic diseases.

In this study, we show that tanycytic ANXA1 contributes to the counterregulatory response to hypoglycemia, promoting an increase in glucose levels. This tanycyte function in glycemic control further supports previous findings showing that tanycyte depletion impairs the counterregulatory response^66^, and that tanycyte-selective leptin receptor deletion disrupts glucose homeostasis^67^. Our study also reveals a novel role for tanycytes in thermoregulation: tanycytic ANXA1 appears to decrease temperature through the sympathetic nervous system and BAT activity, likely via the DMH. The ANXA1/formyl peptide receptor (FPR) pathway has already been implicated in temperature regulation. Notably, exogenous lipocortin (former name for ANXA1) reduces fever induced by cytokines or LPS^68,69^, highlighting its antipyretic properties^70^. Similarly, FPR2 can also decrease temperature by binding ceramide in BAT^71^. Conversely, FPR2 KO mice display an increase in body temperature under high-fat diet, associated with an increase in *Ucp2* expression in muscles^72^. Finally, our study further supports the notion that tanycytes exert bimodal effects on physiological functions: indeed, in this study, both the ANXA1 analog and knockout led to a reduction in food intake. Such dual actions are not unexpected for tanycytes and may reflect their glial nature, which can influence energy balance in both orexigenic and anorexigenic directions and through diverse mechanisms^12^. Additionally, ANXA1 itself is a complex molecule: its N-terminal domain signals through FPR1/2 receptors—associated with reduced food intake through Ac2-26 in our study—while its C-terminal domain has been linked to decreased PGE2 production, an anorexigenic mediator^73,74^. Moreover, ANXA1 may induce both anti- and pro-inflammatory responses depending on several factors such as targeted cell type or post-translational modifications^75^.

Tanycyte ANXA1 highlights a crucial interplay between neuroinflammation and metabolism in the regulation of energy balance, notably to fine-tune adaptive physiological responses. This finding complements previous studies linking tanycytes to inflammatory processes^43^, including inflammation-induced anorexia via NF-κB signaling^73^. The critical role of ANXA1 at the crossroads of metabolism and neuroinflammation becomes especially evident during pathological conditions such as type 2 diabetes and obesity. In humans, plasma ANXA1 levels are elevated in diabetic patients^42^ while findings in obese patients are more variable, with reports of both increased and decreased levels (some studies report increased serum ANXA1 levels^41^, while others observe decreased levels^76^). Despite this, genetic variations in *ANXA1* do not appear to be significantly associated with type 2 diabetes risk^77^. In rodents, total ANXA1-deficient mice exhibit increased susceptibility to diet-induced obesity, insulin resistance, and metabolic dysfunction^36,42^. Conversely, treatment with recombinant ANXA1 improves insulin sensitivity and reduces metabolic complications^42^. Mechanistically, *AnxA1* expression is upregulated during adipogenesis^36,76^ and influences adipocyte function by regulating lipid metabolism and adipokine secretion, thereby impacting insulin sensitivity and energy storage^42^. More precisely, it exerts anti-inflammatory effects by modulating leukocyte migration and cytokine production, mitigating the chronic low-grade inflammation associated with obesity and insulin resistance. In contrast to peripheral tissues, where ANXA1 increases during inflammation, we show that its hypothalamic levels decrease after long-term high-fat feeding—potentially contributing to local neuroinflammation and metabolic dysregulation^43^. Insulin resistance may be a cause of this decrease, although *InsR* depletion in tanycytes does not induce changes in ANXA1 expression^45^. Consistently, tanycyte-specific ANXA1 deletion promotes a trend to gain weight, and elevated body temperature—features that mimic the obese state.

Although our study was conducted in a mouse model, similar mechanisms likely exist in the human brain. Indeed, tanycytes also line the bottom of the third ventricle in humans^17,78^ and express ANXA1, as shown by recent single-nucleus RNA sequencing and spatial transcriptomics studies^79^. Additionally, ANXA1 protein has also been detected in CSF-derived EVs^65^ and in the ependyma^57^. However, while peripheral *ANXA1* levels are known to increase during obesity in humans^41^, its central regulation remains unknown, as current transcriptional datasets are limited to individuals with normal BMI^79^. Measuring ANXA1 levels in CSF could provide valuable insight into its role in the human brain, particularly in the context of metabolic disorders.

In conclusion, this study advances our understanding of tanycyte-mediated communication in the hypothalamus and highlights ANXA1-containing EVs as actors in the control of neuroinflammation, energy balance, and thermogenesis. By uncovering the regulatory mechanisms and physiological impacts of tanycyte-derived EVs, this work opens new therapeutic avenues for addressing metabolic disorders such as obesity and type 2 diabetes. Future research should focus on further elucidating the EV cargo, the signaling pathways it modulates, and how this tanycyte–microglia axis might be harnessed to restore hypothalamic homeostasis.

## RESOURCE AVAILABILITY

### Lead contact

Requests for further information and resources should be directed to and will be fulfilled by the lead contact.

### Materials availability

This study did not generate new unique reagents.

### Data and code availability

All proteomics data have been deposited in the PRIDE repository under accession number XXXX and are publicly available as of the date of publication. This paper does not report original code. Any additional information required to reanalyze the data reported in this paper is available from the lead contact upon request.

## Supporting information

Supplemental Media 1

Supplemental Media 2

## ACKNOWLEDGEMENTS

This work was supported by the European Research Council Starting Grant (TANGO, No. 948196). F.L. is supported by the Swiss National Science Foundation (PCEFP3_194551, 320030-231343), the Novartis Foundation for medical-biological research, and the Department of Biomedical Sciences at the University of Lausanne. Mass spectrometry-based proteomics work was performed by the Protein Analysis Facility of the Faculty of Biology and Medicine, University of Lausanne, Lausanne, Switzerland. The authors also thank the Animal Facility (UNIL), the Histology Facility (HF-UNIL), the Cellular Imaging Facility (CIF-UNIL), and the Electron Microscopy Facility (EMF-UNIL).

## AUTHOR CONTRIBUTION

R.D. designed and performed experiments, analyzed and interpreted the data, produced the figures, and wrote the manuscript. C.G., I.K., T.D., A.R., J.B., and J.E.-M. performed experiments and analyzed data. D.L.-R. analyzed bioinformatic data and produced the figures. E.DiV. designed and produced *Tshr*-Cre viruses. F.L. designed experiments, oversaw research, analyzed and interpreted the data, wrote the manuscript, and produced the figures.

## SUPPLEMENTAL INFORMATION TITLES AND LEGENDS

Document Supplemental Figures. Figures S1-S7

Supplemental Table 1

Supplemental Table 2

Supplemental Table 3

Supplemental Media 1

Supplemental Media 2

**<u>Supplemental Table 1.</u>** Proteomic analysis on tanycyte cells, EV, and soluble secretome fractions. Three technical replicates in 3 cultures were analyzed (n=9 per group).

**<u>Supplemental Table 2.</u>** BW, delta BW, delta BW during fasting, glycemia, core temperature, and corticosterone levels in TanAnxa1^-/-^ mice.

**<u>Supplemental Table 3</u>**. Key resources table.

**<u>Supplemental Media 1.</u>** Video showing a series of images acquired by inverse contrast scanning electron microscopy, illustrating the ultrastructure of multivesicular cargos in adult male mice.

**<u>Supplemental Media 2.</u>** Video showing a series of images acquired by inverse contrast scanning electron microscopy, illustrating the ultrastructure of multivesicular cargos in primary tanycyte culture.

## METHODS

## EXPERIMENTAL MODEL AND STUDY PARTICIPANT DETAILS

### Mice and genotyping of transgenic animals

Two-to-six-month-old male C57BL/6J mice (initially obtained from Charles River), Rosa26-floxed stop tdTomato^53^, Rosa26-floxed stop tdTomato::*Sf1-*CRE mice^80^, *Pomc*-GFP mice^81^, Rosa26-floxed stop tdTomato::B6.Cg-Tg(Camk2a-cre)T29-1Stl/J (The Jackson Laboratory, Cat#005359), and *Anxa1*-floxed mice (GemPharmatech) were used in this study. Animals were housed in groups (from 2 to 5 mice per cage) and maintained in a temperature-controlled room (at 22–23 °C) on a 12 h light/dark cycle with ad libitum access to diet and water. The diets used in this study are a normal diet (Safe, 105 SP 25), a 50%/35% high-fat high-sucrose diet (HFHS diet) (Safe, U8955 Version 9), and a 60% high-fat diet (60HF diet) (Safe, U8954 Version 205). Biopsies were collected for genotyping, and DNA extraction was performed using the HotShot method. PCR amplification was performed using KAPA2G Fast ReadyMix (KAPA Biosystems, KK5103) following the manufacturer’s instructions. The primers (5′-3′) used in this study are available on request. All animal procedures were performed at the University of Lausanne and were reviewed and approved by the Veterinary Office of Canton de Vaud (VD3634, VD3872, and VD3825).

### Primary cell cultures

Hypothalamic median eminences were isolated from 10-day-old (P10) pups (Sprague Dawley rat, Janvier Laboratory) for primary tanycyte cultures, as previously described^82^. Briefly, after decapitation and collection of the brain, MEs were dissected and crushed on 40-µm nylon mesh (VWR, #732-2757). Dissociated cells were cultured in high-glucose tanycyte culture medium (TCM) (DMEM high glucose (Gibco, cat no 419666052), 10% pen-strep, 2% L-glutamine) supplemented with 10% (v/v) donor bovine serum at 37°C under a humid atmosphere of 5% CO2. The culture medium was changed after 5 days of culture and subsequently every 2 days. On reaching confluence, tanycytes were seeded in 24-well plates on poly-L-lysine-coated glass coverslips for immunocytochemistry, uncoated plastic 12-well plates for gene expression, and uncoated plastic 6-well plates for protein expression analysis. For EV production, primary tanycytes were seeded in T175 flasks.

Mouse brains were collected from P3-P5 pups for primary microglia culture, as previously described^83^. Briefly, mice were sacrificed by decapitation, and the brain was collected in HBSS. The olfactory bulb and cerebellum were removed, and the meninges were peeled off. The tissue was grossly minced, transferred into TripLE Express Enzyme Solution (cat no 12-604-021, Life Technologies), and incubated at 37°C for 20min followed by mechanical dissociation. DMEM high glucose (cat no 419666052, Gibco) supplemented with 10% fetal bovine serum and 1% pen-strep was added to the preparation at a 3:1 ratio. Cells were pelleted at 400x g for 4 min at RT. The pellet was resuspended in a fresh culture medium and plated in a T75 cell culture flask. The mixed astrocyte/microglia cultures were maintained at 37°C and 5% CO2. After two weeks, microglia were detached from the underlying astrocyte layer by smacking the flask. Detached microglia contained in the astrocyte-conditioned medium (ACM) were then pelleted at 400 g for 4 min, resuspended in the ACM, and seeded on poly-D-lysine-coated plates.

### rAAV vectors production

Recombinant adeno-associated viral vectors (rAAV) were produced by the GIGA-Viral Vector platform. Briefly, pAAV Tshr mCherry P2A Cre (Vector Builder # VB220222-1152gmq) or pAAV CAG mcherry P2A CRE (Vector Builder # VB220223-1220nbv) plasmids were co-transfected into 293AAV Cell Line (Cell Biolabs, AAV-100) together with a helper plasmid (Part No. 340202 VPK-401 kit) and REP-Cap plasmid (mix of various serotypes at ratio 1:1: pAAV-1 [Cell Biolabs #VPK-421] and pAAV-2 [Cell Biolabs #VPK-422]. After collecting the cell supernatant, rAAV vectors were concentrated and titrated using ABM good kit (#GE931) at a concentration of >1E+13 genome copy/mL (GC/mL).

## METHOD DETAILS

### Stereotactic surgery on mice

To recombine DNA in tanycytes, TAT-CRE fusion protein (MERCK, SCR508; diluted by 1:4 in saline solution; 1 μl over 3 minutes) or AAV-Tshr-Cre (2.77E+13 TU/mL; 1 μl over 3 minutes) was stereotactically infused into the dorsal third ventricle (AP= 0 mm and ML= 0 mm from the bregma; DV= –3.9 mm from the cortex surface) or the lateral ventricle (AP= −0,3 mm and ML= 1 mm from the bregma; DV= –2.5 mm from the cortex surface) of ketamine/xylazine-(100 mg/kg and 20 mg/kg, respectively) or isoflurane-anesthetized mice, as previously described^53,66^. For *Anxa1*-flox mice, control (+/+), heterozygous (flox/+), and homozygous (flox/flox) littermate mice were infused with AAV-*Tshr*-Cre. The efficiency and specificity of DNA recombination were validated by *in situ* hybridization, immunohistochemistry, or/and qPCR.

For Ac2-26 infusion, mice were canulated in the lateral ventricle a week before the experiment at the coordinates from the bregma of AP= −0.3 mm; ML= −1 mm; DV= −2.3 mm (from cortex surface) and infused with Ac2-26 or Scrambled peptide (Isca Biochemicals; PP-100 and PP-100 SCRAMBLED (design based on Kamal et al., 2001); 1ug, 2ug or 4ug; 1ul over 1 minute) under isoflurane anesthesia.

### Mouse physiology

For energy balance analysis, body and food weight were measured using a scale. Body composition was recorded using nuclear magnetic resonance (EchoMRI™). Blood glucose measurements were made from tail vein blood using a Contour®NEXT glucose monitor. Body temperature was recorded using a rectal thermometer (BIOSEB, BIO-TK8851). For BAT temperature, mouse backs were shaved the day before the experiment, and the temperature was recorded using a thermal camera (FLIR, E53-24).

For the fasting-refeeding paradigm, mice were fasted for 24 hours, then isolated 1 hour before the experiment (starting at 9 a.m.), and food weight, glycemia, and body temperature were measured at different time points during refeeding. For the 2-deoxyglucose (2-DG, Sigma-Aldrich, D8375) test, 2-DG was injected i.p. at 500mg/kg in saline, and glycemia and body temperature were measured at different time points. For the norepinephrine (NE, Sigma-Aldrich, A0937) test, NE was injected i.p. at 1mg/kg in saline, and glycemia and body temperature were measured at different time points. For the β3 receptor antagonist SR59230A (B3RA, Sigma-Aldrich, S8688) test, B3RA was injected i.p. at 3mg/kg in saline, and glycemia and body temperature were measured at different time points. For cold exposure, mice were exposed at 4°C for 4 hours: glycemia and body temperature were measured at different time points.

### Metabolic cages

Total energy expenditure, oxygen consumption, carbon dioxide production, food intake, and ambulatory movements were measured using calorimetric cages (SABLE, Promethion systems) for up to 10 days. Mice were individually housed and acclimatized to the cages for 48 hours before experimental measurements. For the fasting-refeeding paradigm, mice were fasted for 24 hours and then refed. For Ac2-26 injection, mice received the i.c.v. injection through the cannula after 24 hours of fasting. For the thermoneutrality experiment, mice were exposed at 30°C. For cold exposure, mice were exposed at 4°C.

### Energy imbalance paradigm for tissue collection

Mice were killed in different metabolic states, and tissues were collected for further analyses. The different time points used for the fast-refeeding paradigm were fed, 24h-fast, 2h-refed, and 4h-refed. Fed mice were killed at 8 a.m. Fasted mice were fasted starting at 8 a.m. and had access to water. Refed mice were fasted for 24 hours and received *ad libitum* diet access to food beginning at 8 a.m. Body weight and glycemia were measured at each time point to control the metabolic state.

Three different paradigms were used for the high-fat diet: the “short-term” 60HF and HFHS diets refer to 1h, 3h, and 6h food exposure. Food was removed from cages at 4 p.m. and 60HF or HFHS diets were introduced at 6 p.m. for 1 to 6 h (all mice were exposed to 60HF and HFHS diets for 10 minutes the day before for habituation). The “middle-term” 60HF and HFHS diets refer to 3 days, 14 days, and 28 days of food exposure. The “long-term” 60HF and HFHS diets refer to 8 weeks of food exposure.

To test the impact of ANXA1, scrambled or Ac2-26 peptide was infused (i.c.v., 1ug, 2ug, or 4ug) in fed or 24h-fasted mice 2h before sacrifice.

### Tissue collection

For endocrine parameters, blood was collected by cardiac puncture after anesthesia with pentobarbital using a syringe containing EDTA.

For immunohistochemistry and *in situ* hybridization on fixed tissue, mice were anesthetized with pentobarbital and perfused transcardially with 0.9% saline, followed by an ice-cold solution of 4% paraformaldehyde in 0.1 M phosphate buffer (pH 7.4). Brains were quickly removed, postfixed in the same fixative for 2h at 4°C, and immersed in 20% sucrose in 0.1M phosphate-buffered saline (PBS) at 4°C overnight. Alternatively, while collecting multiple organs, brains were quickly removed and fixed by immersion in 4% paraformaldehyde in 0.1 M phosphate buffer (pH 7.4) at 4°C overnight. Brains were finally embedded in ice-cold OCT medium (optimal cutting temperature embedding medium, Tissue Tek, Sakura) and frozen on dry ice or in liquid nitrogen-cooled isopentane.

For electron microscopy, mice were anesthetized with isoflurane, and perfused transcardially with 0.9% saline followed by an ice-cold solution of 2% paraformaldehyde/2% glutaraldehyde in 0.1 M phosphate buffer, pH 7.4. Brains were quickly removed, postfixed in the same fixative overnight at 4°C. Two-hundred-micrometer-thick hypothalamic slices were then cut using a vibratome. TdTomato fluorescence in tanycytes was then acquired using a ZEISS Axio Imager.M2 microscope equipped with ApoTome.2 to give coordinates to each protrusion in the slice. Afterwards, the samples were incubated in 2% (wt/vol) osmium tetroxide and 1.5% (wt/vol) K4[Fe(CN)6] in 0.1 M PB buffer for 1 hr, followed by one-hour incubation in 1% (wt/vol) tannic acid in 0.1 M PB buffer. Subsequently, brain slices were incubated in 1% (wt/vol) uranyl acetate for 1 hr and dehydrated at the end of standard gradual dehydration cycles in ethanol. Samples were flat-embedded in Epon-Araldite mix^84,85^. All procedures were performed at room temperature.

For brown adipose tissue (BAT) histochemistry and qPCR, mice were anesthetized with pentobarbital and sacrificed by decapitation. Interscapular BAT was dissected, with one half frozen in liquid nitrogen for qPCR analysis and the other half fixed in 10% neutral buffered formalin (Sigma-Aldrich, #HT501128) for 24 hours at room temperature for histological analysis. The fixed tissue was then embedded in paraffin, sectioned, and stained with hematoxylin and eosin.

### Primary culture treatment for sample collection

Before the experimental assays described below, tanycytes were washed twice with PBS and cultured without serum for 24h.

To test the impact of glucose, primary tanycytes were incubated in no-glucose tanycyte culture medium (TCM) (DMEM no glucose (Gibco, P04-01548S1), 10% pen-strep, 2% L-glutamine) for 24h. The medium was then changed for high-glucose TCM before cell harvesting at different time points. Alternatively, primary tanycytes were incubated in no-glucose TCM supplemented with 0.5mM glucose.

To test the impact of hormones, primary tanycytes were incubated with 8-CPT-cAMP (100 μM), dexamethasone (1 μM), insulin (100nM), or vehicles in high-glucose TCM before cell harvesting at different time points.

To test the impact of calcium signaling, primary tanycytes incubated in no-glucose TCM for 24h were incubated with vehicle (DMSO) or BAPTA (40mM) 30 min before being exposed to high-glucose TCM, still in the presence of vehicle or BAPTA: cells were then harvested at different time points. Alternatively, primary tanycytes were incubated in a glucose-free Kreb solution (in mM: NaCl 135.5, MgCl_2_ 1.2, KCl 5.9, HEPEs 11.5, CaCl_2_ 1.8, final pH 7.3) for 24h, then with glucose Kreb solution (glucose 11.5mM) or a Ca^2+^-free glucose Kreb solution (in mM: NaCl 135.5, MgCl_2_ 1.2, KCl 5.9, glucose 11.5, HEPES 11.5, 200 μM Na-EGTA, final pH 7.3): cells were harvested at different time points.

To test the impact of calcium stores, tanycytes were incubated in Ca^2+^-free glucose Kreb solution for 24h and then treated with 1 µM thapsigargin (or DMSO) 15 min before cell harvesting.

Microglia were cultured in DMEM high glucose supplemented with 1% pen-strep, without serum, for 24h. They were then treated with 5uM Ac2-26 or scrambled peptide for 2h and 4h. For synaptosome uptake assays, synaptosomes were isolated from dissected hypothalami using the Syn-PER reagent (Thermofischer, #87793) following the manufacturer’s instructions. Microglia were cultured with 5 µM Ac2-26 or scrambled for 1h before incubation in a medium containing synaptosomes with 5 µM Ac2-26 or scrambled. Cells were harvested 2h later.

### Sample collection from cell culture

Extracellular vesicles were isolated by ultra-centrifugation, as described previously (Lasser et al., 2012). Briefly, the culture media (depleted from serum-EVs) of primary tanycytes were collected and centrifuged at 300 x g for 5 min to pellet cells and then at 2000 x g for 10 min to discard dead cells. Supernatants were centrifuged at 10 000 x g for 30min to remove cell debris. EVs were then isolated from the soluble secretome by ultracentrifugation at 100 000 x g for 2 hours. The pellet containing extracellular vesicles was washed with PBS and re-centrifuged at 100 000 x g for 2 hours. Extracellular vesicles were collected in a minimal volume of PBS. Extracellular vesicle size distribution was measured by Nanoparticle Tracking Analysis using the NanoSight system (NanoSight, UK). The EV pellet was resuspended in 100 μL lysis buffer (150M NaCl, 1% NP-40, 50mM Tris-HCl pH 8.0, 0.5% Sodium deoxycholate, 0.1% SDS, Protease inhibitor, PhosphoStop) and protein concentration was determined by Bradford assay (23227, Thermo Fisher).

For immunocytochemistry, cells were washed three times with PBS and fixed in 4% PFA for 5 minutes at room temperature. After fixation, cells were washed with PBS and immediately processed for immunocytochemistry.

For electron microscopy, cells were fixed with glutaraldehyde 5% in phosphate buffer 0.2M, pH 7.4, added to the equal amount of cells’ growth media. for 15 min, and then glutaraldehyde 2.5% in phosphate buffer 0.1M for 1h at RT. Following the fixation, the samples were incubated with 1% osmium tetroxide for 1 hour on ice. After the extensive wash cycles with distilled water, a subsequent 1-hour incubation with 1% uranyl acetate was performed. Dehydrated samples were embedded in a drop of Epon-Araldite mix (EMS) directly added to the glass slide surface and polymerized for 24 hours at 60 °C. Samples were re-embedded on support and removed by the hot-cold shock procedure before the sectioning step.

For Western blot, cells were washed twice with ice-cold PBS, before adding 100ul of lysis buffer (150M NaCl, 1% NP-40, 50mM Tris-HCl pH 8.0, 0.5% Sodium deoxycholate, 0.1% SDS, Protease inhibitor, PhosphoStop). Cells were scraped and processed for protein quantification by the BCA protein assay kit (23227, Thermo Fisher).

For qPCR, cells were washed twice with ice-cold PBS before adding 350 μL of lysis buffer (RNeasy Mini Kit, #74104, Qiagen). Cells were scraped and collected into an Eppendorf tube for homogenization.

### Proteomics analysis

Cell pellets were digested following a modified version of the iST method^86^ (named miST method). Washed cell pellets were lysed in 100 μL miST lysis buffer (1% Sodium deoxycholate, 100mM Tris pH 8.6, 10 mM DTT) and heated for 10min at 75 °C. Based on tryptophan fluorescence quantification^87^, 30 μg of proteins at 1 ug/ul in miST lysis buffer (1% Sodium deoxycholate, 100mM Tris pH 8.6, 10 mM DTT), were transferred to new tubes. Samples were heated for 5 min at 95°C, diluted 1:1 (v:v) with water containing 4mM MgCl2 and benzonase (Merck #70746, 100x dil of stock = 250 Units/ul), and incubated for 15 minutes at RT to digest nucleic acids. Reduced disulfides were alkylated by adding ¼ vol. of 160 mM chloroacetamide (32 mM final) and incubating for 45min at RT in the dark. Samples were adjusted to 3 mM EDTA and digested with 0.5 μg Trypsin/LysC mix (Promega #V5073) for 1h at 37°C, followed by a second 1h digestion with an additional 0.5 μg of proteases. To remove sodium deoxycholate, two sample volumes of isopropanol containing 1% TFA were added to the digests, and the samples were desalted on a strong cation exchange (SCX) plate (Oasis MCX; Waters Corp., Milford, MA) by centrifugation. After washing with isopropanol/1%TFA, peptides were eluted in 200 μL of 40% MeCN, 19% water, 1% (v/v) ammonia, and dried by centrifugal evaporation. Pelleted vesicles in PBS were supplemented with concentrated sodium deoxycholate (DOC) and DTT to reach a final 1% DOC and 3mM DTT in 300 μL. Samples were processed as described above, with the difference that no benzonase treatment was performed. Soluble secretome samples in ammonium bicarbonate buffer were lyophilized and resuspended in 50 μL miST buffer and digested as described above for the cell pellets, but without benzonase treatment.

Liquid Chromatography-Mass Spectrometry (LC-MS/MS) analyses were carried out on a TIMS-TOF Pro (Bruker, Bremen, Germany) mass spectrometer interfaced through a nanospray ion source (“captive spray”) to an EvoSep One liquid chromatography system (EvoSep, Odense, Denmark). Peptides were resuspended in 0.1% formic acid and separated on a reversed-phase 15 cm C18 column (150 μm ID, 1.5 μM, EvoSep Product EV1137) at a flow rate of 220 nl/min with a standard EvoSep 15 sample per day method (runtime time: 88 min, solvents were water and acetonitrile with 0.1% formic acid, useful gradient 0-35%). Data-independent acquisition was carried out using a method similar to a standard DIA-PASEF method reported previously^88^, with ion accumulation for 100 ms for each of the survey MS1 scan and the MS2 scans. Duty cycle was kept at 100%. Precursor ions were chosen within a reduced mobility range from 1/k0 =0.8 to 1.3 and between m/z=400 to 1200. Collision energy was ramped linearly based uniquely on the 1/k0 values from 20 (at 1/k0=0.6) to 59 eV (at 1/k0=1.6). Per cycle, the mass range 400-1200 m/z was covered by a total of 32 windows, each 25 Th wide. Two windows were acquired per TIMS scan (100ms) so that the total cycle time was 1.8s.

Identification of peptides directly from DIA data was performed with Spectronaut 19.7 with the Pulsar engine and searching the reference Rattus norvegicus proteome (www.uniprot.org) database, version of February 14th, 2024 (47’943 sequences), and a contaminant database containing the most usual environmental contaminants and enzymes used for digestion^89^. For identification, peptides of 7-52 AA length were considered, cleaved with trypsin/P specificity and a maximum of 2 missed cleavages.

Carbamidomethylation of cysteine (fixed), methionine oxidation, and N-terminal protein acetylation (variable) were the modifications applied. FDRs for peptide and protein group identifications were all at 1%. Ion mobility for peptides was predicted using a deep neural network and used in scoring. The libraries created contained overall 141’934, 88’603, and 50’101 precursors for cell pellets, EV, and soluble secretome, respectively.

Peptide-centric analysis of DIA data was done with Spectronaut 19.7 using the library generated by Pulsar from DIA data. Single-hit proteins (defined as matched by one stripped sequence only) were kept in the Spectronaut analysis. Peptide quantitation was based on XIC area, for which a minimum of 1 and a maximum of 3 (the 3 best) precursors were considered for each peptide, from which the median value was selected. Quantities for protein groups were derived from inter-run peptide ratios based on the MaxLFQ algorithm^90^. Data for cell pellets, EV, and soluble secretome samples were analyzed separately with Spectronaut. Normalization by the software was applied for cell pellets but not for EV and soluble secretome data. When done, normalization was based on the median of peptides. Overall, 133’709, 84’265, and 48’192 precursors were quantified for cell pellets, EV, and soluble secretome, respectively. These mapped, respectively, to 8’338, 3’515, and 4’365 protein groups. The average number of data points per LC peak was between 6.3 and 7.7.

All subsequent data analyses were done with an in-house developed software tool (available on https://github.com/UNIL-PAF/taram-backend). Contaminant proteins were removed, and quantity values for protein groups generated by Spectronaut were log2-transformed. After assignment to groups, only proteins quantified in at least 3/3 samples in at least one group were kept. Missing values were imputed based on a normal distribution with a width of 0.3 standard deviations (SD), down-shifted by 1.8 SD relative to the median. Student’s T-tests were carried out among conditions, with Benjamini-Hochberg correction for multiple testing (adjusted p-value threshold <0.05). Imputed values were later removed. The differences of means obtained from the tests were used for 1D enrichment analysis on associated GO/KEGG annotations, as described^91^. The enrichment analysis was also FDR-filtered (Benjamini-Hochberg, adjusted p-value <0.02).

### Molecular biology

For qPCR, the RNeasy Mini kit (#74104 QIAGEN) was used to extract mRNA from frozen tissues and cultured cell samples. Approximately 250 ng of RNA for tissue and 100ng for cell culture was then reverse-transcribed using the M-MLV reverse transcriptase (M3683, Promega) following the manufacturer’s instructions. cDNAs were diluted (1:10), and qPCR was performed using GoTaq qPCR Master Mix (Promega). The primers (5′-3′) used in this study are available on demand.

For Western Blot, 5ug of proteins were loaded using Laemmli loading buffer in a 15% Acrylamide/Bis Gel (Tris pH8.8 1.5mol/l, acrylamide/bis 15%, SDS 20%, APS 10%, Temed), migrated for 40min at 90V and then 1h at 120V, and then transferred to a PVDF membrane for 1h at 120V. For extracellular vesicle extracts, proteins were migrated on Mini-PROTEAN TGX (Biorad, #4561084) SDS-PAGE gels. Immunolabelling of the membrane was then performed using primary antibody overnight in 5% BSA 1x TBS-Tween 0.1% at 4°Cand then with secondary antibodies (1:15’000 in 1x TBS 0.1% Tween). The revelation was done using the LiCor Odyssey Fc. The antibodies and dilutions used in this study are reported in Supplemental Table 3.

### Immunohistochemistry and Immunocytochemistry

Brains were cut using a cryostat into 20 μm thick coronal sections (or 25 μm thick coronal sections for c-Fos and pSTAT3 immunostaining) and processed for immunohistochemistry as described previously. The antibodies used in this study are reported in Supplemental Table 3. For most of the antibodies, slide-mounted sections were: 1) blocked for 1h using a solution containing 4% normal goat serum and 0.3% Triton X-100, 2) incubated overnight at 4°C with primary antibodies followed by 2h at room temperature with a cocktail of secondary Alexa Fluor-conjugated antibodies (1:500, Molecular Probes, Invitrogen, San Diego, CA, Supplemental Table 3) and 3) counterstained with DAPI (1:10,000, Molecular Probes, Invitrogen), and 4) coverslipped using Mowiol (Calbiochem, La Jolla, CA). For c-Fos and pSTAT3 immunostaining, slices were permeabilized for 30min at RT in 1% Triton X100 and blocked in a 4% BSA and 0,3% Triton X-100 in PBS 1X blocking solution for 1h at 37°C. The same protocol was used for cell cultures.

### In situ hybridization

Fixed frozen brains were cut using a cryostat into 20-μm-thick coronal sections and processed for RNAscope® *in situ* hybridization following the manufacturer’s instructions (ACD). Slide-mounted sections were first 1) posted fixed in 4% PFA in 1X PBS for 15min at 4 degrees 2) dehydrated by successive incubation in 50%, 70%, and 100% EtOH for 5 min each and air dried for 5min 3) incubated in a boiling 1X Target retrieval solution for 5 minutes and 4) incubated at 40°C with Protease III solution for 30 minutes. The sequential hybridizations were then performed following the manufacturer’s instructions. The probes used in this study are reported in Supplemental Table 3.

### Fluorescent Microscope

Images were acquired with either a Leica Thunder Imaging System, an i90 Nikon microscope, or a confocal (Zeiss LSM 780, 800, and 900). Frames were collected stepwise over a defined z-focus range corresponding to all visible fluorescence within the section: multiple-plane frames were collected at a step of 1 µm while using x20, x40, or x63 objective (between 4 and 10 frames per image). All images were then saved in .lif or .nds, processed to get orthogonal and maximal intensity projections, and finally exported in .tiff for the processing steps (i.e., adjust brightness and contrast, change colors, and merge images using ImageJ). The original DAPI blue pseudo color was changed to white.

### Electron microscopy^92,93^

For the tissue samples, flat-embedded samples were mapped using binoculars to map the precise region of interest at the basal part of the ventricle. For the ultramicrotome sectioning, samples were carefully oriented inside the holder according to the correlation maps derived from the embedded samples. Polymerized flat blocks were oriented perpendicular to the vibratome section plane and trimmed using a 90° diamond trim tool (Diatome, Biel, Switzerland) to delimit the area of interest.

Samples from the cell culture were sectioned in a frontal view, after the observation under the microscope, to detect the ROI populated with cells.

The arrays’ sequential sections of 100 nm were generated using a 35° ATC diamond knife (Diatome, Biel, Switzerland) mounted on a Leica UC6 microtome (Leica, Vienna). Arrays that covered the span of the ROI were transferred to 2×4 cm pieces of silicon wafer ^94^ and analyzed using FEI Helios Nanolab 650 scanning electron microscope (Thermo Fischer, Eindhoven). Wafers were screened to target the relevant sections, and the ventricle area was imaged by tiling multiple images using the following imaging settings: M.D. detector, accelerating voltage 2kV, current 0.8nA, dwell time 4-6 µs with images collected in the inverted mode. Subsequently, relevant areas were re-imaged using higher resolution parameters. with overall resolution ranging between 2 and 10 nm pixel size. Images were collected either manually or using the AT module of the MAPs program (Thermo Fischer, Eindhoven)^94^.

Modeling and visualization were performed using 3dmod. For electron microscopy data interpretation, previous reports in the literature were used to recognize the different neural cell types based on their ultrastructural characteristics^16,95^. Stitched sequential images were aligned into a stack using AMIRA (Thermo Fischer Scientific) or the Fiji TrakEM module (Ref). These stacks of images were manually traced, following the contours of different structures (cell membranes, organelles, and EVs), using the 3Dmod module of the IMOD program. The traced outlines were transformed into the meshed skins used to model the interactions in the movies (Ref Mastronarde).

## QUANTIFICATION AND STATISTICAL ANALYSIS

### Proteomics analysis

Downstream proteomics data analysis of the soluble secretome, extracellular vesicles, and cell was conducted in R (v4.5.0) using the protti package (v0.9.1). To assess data variance, principal component analysis (PCA) was performed, followed by differential expression analysis using multiple t-tests with Benjamini–Hochberg (BH) correction. Functional enrichment analysis of differentially expressed proteins or the most abundant proteins per condition was carried out using the enrichR (3.4) package, querying the “GO_Biological_Process_2023”, “GO_Molecular_Function_2023”, and “GO_Cellular_Component_2023” libraries.

### scRNAseq data analysis

The publicly available scRNA-seq datasets Brunner et al. (2024) and Steuernagel et al. (2022) were downloaded from GEO (accession: GSE266664) and CellxGene, respectively. Both datasets were obtained as pre-processed Seurat objects, in which quality control, normalization, dimensionality reduction, and cell type annotation had already been performed as described in the respective publications. Feature expression plots were generated using the FeaturePlot() function in Seurat (v5.2.0), based on the original HypoMap embeddings and cell type classifications. Functional enrichment analysis was conducted using the GSVA package (v22.0) with the ssGSEA method (Barbie et al., 2009). ssGSEA is a single-sample, non-parametric gene set enrichment method that calculates the normalized difference between the empirical cumulative distribution functions of gene expression ranks inside and outside a given gene set. Feature co-expression plots were generated using ggplot2 (v) after fetching data from the Seurat objects using the FetchData() function. All analyses were performed in R (v4.5.0).

### Image analyses

For brain image analyses, the entire mediobasal hypothalamus was divided into four subregions on the anteroposterior axis, corresponding to zone 1 (from bregma −1.3 to −1.6mm), zone 2 (from bregma −1.6 to −1.8 mm), zone 3 (from bregma −1.8 to −2.1 mm), and zone 4 (from bregma −2.1 to - 2.5 mm). These subregions have been previously characterized ^16^. All quantifications were performed using ImageJ (Fiji, Version 2.0.0-rc-69/1.52p).

For the ANXA1 protein, x20 images were acquired along the anteroposterior axis, and the different hypothalamic nuclei were identified using DAPI. Images were processed using “Z-project”, “8-bit conversion”, “set threshold”, and “make binary”. On the composite image, the tanycytes layer facing the vmARH, dmARH, VMH, and DMH was delimited using DAPI (ROI). The Integrated Density was calculated and normalized over the ROI area. ANXA1-positive processes projecting in the VMH and DMH (delimited with DAPI) were quantified manually on 40x images. These analyses were performed on 8 slices per brain (2 per zone).

For *Anxa1* mRNAs, the tanycytes layer facing the vmARH, dmARH, VMH, and DMH was delimited (ROI) using DAPI. *Anxa1* mRNA puncta were manually quantified and normalized over the area. These analyses were performed on 4 slices per brain (1 per zone).

For c-Fos and pSTAT3 activation, x20 images were acquired along the anteroposterior axis, and the different hypothalamic nuclei were identified using DAPI. The number of pSTAT3-or c-Fos-positive cells was quantified on 8 slices per brain (2 per zone) using ImageJ software.

The density of pre-synaptic terminals in hypothalamic nuclei was assessed with VGLUT2 labeling along the anteroposterior and dorsoventral axes. 63x images were processed using “Z-project” and “8-bit conversion”. The number of pre-synaptic puncta was measured using the plugin Puncta Analyzer and normalized over the nucleus area delimited by DAPI.

Microglia and astrocyte density and morphology were assessed using Iba1 and GFAP labeling. Briefly, images were acquired at 20x and 63x along the anteroposterior and dorsoventral axes in the different hypothalamic nuclei delimited using DAPI. They were processed using “Z-project”, “8-bit conversion”, “set threshold”, and “make binary”. Microglia and astrocyte densities were assessed using 20x images: the cell numbers (Iba1/DAPI, GFAP/DAPI) were counted manually and normalized over the nuclei area delimited by DAPI. These analyses were performed on 4 slices per brain (1 per zone). Microglia and astrocyte morphology were assessed using 63x images: ramification was evaluated using the plugin Concentric Circles for the Sholl analysis. Briefly, the cell body was identified as the center of the first circle, and then 6 circles were drawn on the cell with a 2 μm interval between circles. The number of intersections between microglia and astrocyte processes was then counted manually. These analyses were performed on 7 to 10 cells per brain.

For the FPR1 and FPR2 proteins, x20 images were acquired along the anteroposterior axis, and the different hypothalamic nuclei were identified using DAPI. Images were processed using “Z-project”, “8-bit conversion”, “set threshold”, and “make binary”. On the composite image, ARH, ARH, VMH, and DMH were delimited using DAPI (ROI): the Integrated Density was calculated and normalized over the ROI area. These analyses were performed on 8 slices per brain (2 per zone).

For cell culture image analyses, quantifications were performed on multiple tanycytes in at least 3 technical replicates in at least one culture. All quantifications were performed using ImageJ (Fiji, Version 2.0.0-rc-69/1.52p).

Colocalization of ANXA1 and DAPI, CD9, EEA1, lysotracker, and WGA was assessed using Fiji (Version 2.0.0-rc-69/1.52p). In brief, the images were pre-processed using “8-bit conversion”, “Subtract Background”, “Brightness/Contrast”, “Set threshold”, and converted to mask. Colocalization of the masks of the two channels was assessed using “colocalization.”

For tanycyte morphology, x20 images were acquired on bright field. Cell body area and process length were measured using ImageJ.

Microglia CD68-positive phagolysosomal content and synaptosome engulfment were quantified on 63x images. In brief, images were processed using “8-bit conversion”, “Subtract Background”, “Brightness/Contrast”, “Set threshold”, and converted to mask. The percentage of area covered by CD68-positive phagolysosome or synaptosome was assessed using “Measure”.

For BAT images, x20 images were acquired on bright field. Images were then binarized and skeletonized using ImageJ software, and the integrated density was then calculated. Two sections per animal were used for quantification.

### qPCR analyses

Gene expression levels were normalized to actin (*Act*) using the 2−ΔΔCt method and were presented as relative transcript levels.

### Western blot analyses

Densitometric analysis was performed using Image Studio Lite (Version 5.2). B-Actin or revert was used as a control.

### Statistical analysis

All values are expressed as means ± SEM. Data were analyzed for statistical significance with Graph Prism 5 software (Version 11.0), using unpaired t-test, one-way ANOVA followed by a Tukey’s post-hoc test, or two-way ANOVA followed by a Bonferroni’s post-hoc test when appropriate. P-values of less than 0.05 were considered to be statistically significant. Statistics are presented in the figures. The statistical tests used, and the numbers of n, cohorts, or cultures are indicated in the legends.

## ADDITIONAL RESOURCES

All proteomics data have been deposited in the PRIDE repository under accession number XXXX.

## KEY RESOURCES TABLE

Available as Supplemental Table 3

---

**Supplemental Figure 1.**
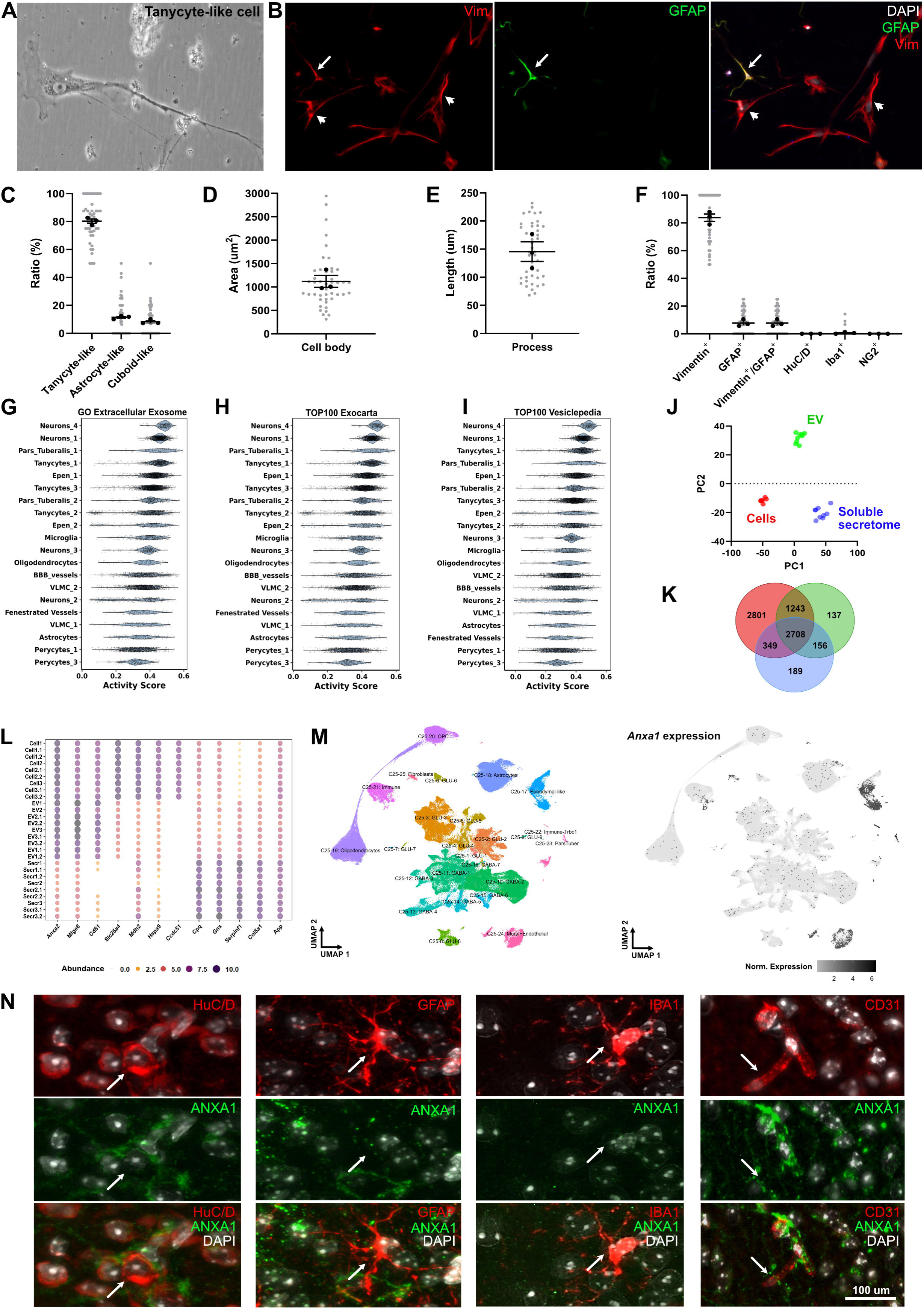
A*n*xa1 expression in the hypothalamus and other brain regions. **(A)** Representative bright-field microscope images of primary tanycyte culture showing elongated and cuboidal cells. **(B)** Representative immunofluorescence of vimentin-positive, GFAP-positive, and both vimentin- and GFAP-positive cells in primary tanycyte culture. Arrows indicate GFAP^+^/Vimentin^+^ cells; Arrowheads indicate GFAP^-^/Vimentin^+^ cells. **(C-F)** Morphological and molecular characterization of primary tanycyte cultures, indicating the cell shape (C), the cell area (D), the process length (E), and the marker distribution (F). 15 pictures per well were analyzed in three technical replicates in one culture (n=3). **(G)** GO_extracellular_exosomes in different cell type clusters from the Brunner/Lopez dataset^20^. **(H-I)** Gene set enrichment analysis on the Brunner/Lopez dataset using Top 100 exocarta (H) and vesiclepedia (I) genes enriched in extracellular vesicles. **(J)** Principal component analysis plot representing proteomics data. **(K-L)** Venn diagram presenting the number of detected proteins (K) and dotplot with selected top markers in each fraction (i.e., cell lysate, EV, and soluble secretome fractions) (L). **(M)** *Anxa1* expression in the mouse hypothalamus using single-cell RNA sequencing (Mouse HypoMap^34^). **(N)** Representative images of ANXA1 staining with HuC/D, GFAP, IBA1, or CD31 staining in the mediobasal hypothalamus in fed male mice. *p < 0.05, \*\**p* < 0.01, \*\*\**p* < 0.001, \*\*\*\**p* < 0.0001.

**Supplemental Figure 2.**
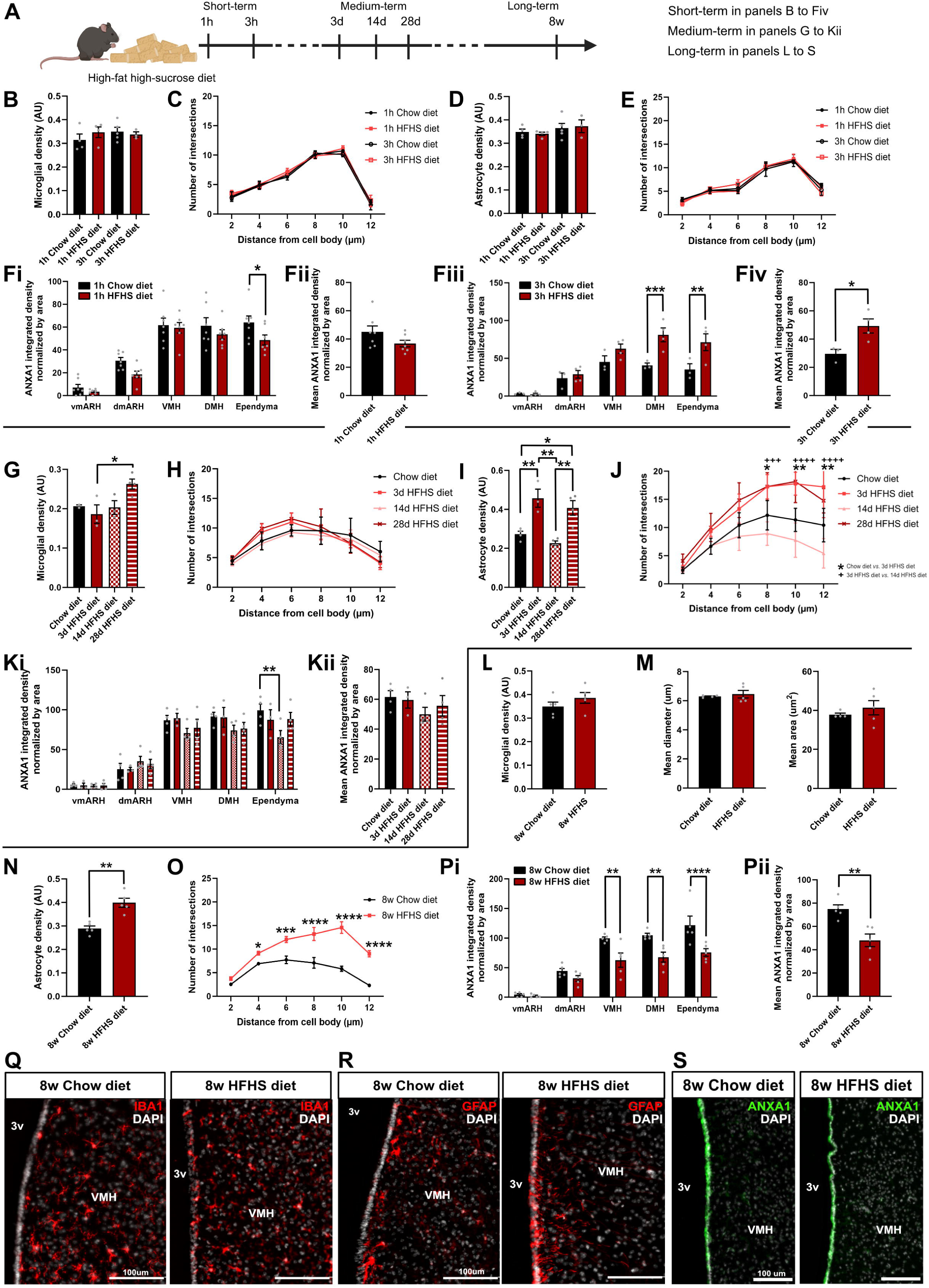
ANXA1 expression in the hypothalamus during high-fat high-sucrose diet paradigms. **(A)** Schematic representations of different high-fat high-sucrose diet paradigms. **(B-F)** Quantifications of microglia density (B) and ramification (C), astrocyte density (D) and ramification (E), and ANXA1 expression along the 3v (F) after short-term HFHS diet (n=3-7 mice/group in two cohorts). Quantifications were performed on the anteroposterior axis (Fi, Fiii) and the entire region (Fii, Fiv). **(G-K)** Quantifications of microglia density (G) and ramification (H), astrocyte density (I) and ramification (J), and ANXA1 expression along the 3v (K) after medium-term HFHS diet (n=3-4 mice/group in one cohort). Quantifications were performed on the anteroposterior axis (Ki) and the entire region (Kii). **(L-P)** Quantifications of microglia density (L), size (M), astrocyte density (N) and ramification (O), and ANXA1 expression along the 3v (P) after long-term HFHS diet (n=4-10 mice/group in two cohorts). Quantifications were performed on the anteroposterior axis (Pi) and the entire region (Pii). **(Q-S)** Representative images of microglia (Q), astrocytes (R), and ANXA1 expression (S) after 8 weeks of HFHS diet. *p < 0.05, \*\**p* < 0.01, \*\*\**p* < 0.001, \*\*\*\**p* < 0.0001.

**Supplemental Figure 3.**
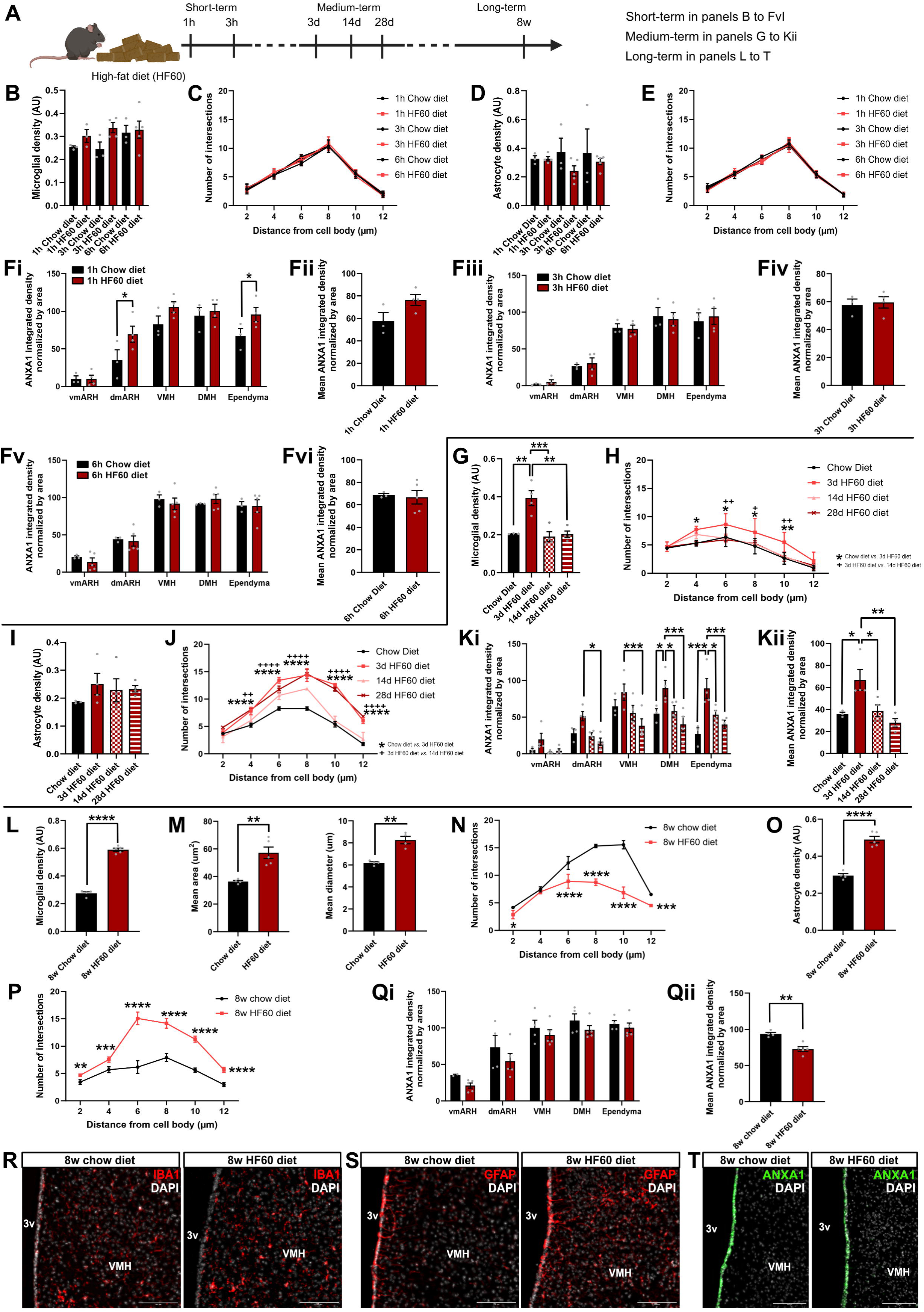
ANXA1 expression in the hypothalamus during high-fat diet paradigms. **(A)** Schematic representations of different high-fat diet paradigms **(B-F)** Quantifications of microglia density (B) and ramification (C), astrocyte density (D) and ramification (E), and ANXA1 expression along the 3v (F) after short-term HF60 diet (n=3-5 mice/group in one cohort). Quantifications were performed on the anteroposterior axis (Fi, Fiii, Fv) and the entire region (Fii, Fiv, Fvi). **(G-K)** Quantifications of microglia density (G) and ramification (H), astrocyte density (I) and ramification (J), and ANXA1 expression along the 3v (K) after medium-term HF60 diet (n=3-4 mice/group in one cohort). Quantifications were performed on the anteroposterior axis (Ki) and the entire region (Kii). **(L-Q)** Quantifications of microglia density (L), size (M), and ramification (N), astrocyte density (O) and ramification (P), and ANXA1 expression along the 3v (Q) after long-term HF60 diet (n=4-5 mice/group in one cohort). Quantifications were performed on the anteroposterior axis (Qi) and the entire region (Qii). **(R-T)** Representative images of microglia (R), astrocytes (S), and ANXA1 expression (T) after 8 weeks of HF60 diet. *p < 0.05, \*\**p* < 0.01, \*\*\**p* < 0.001, \*\*\*\**p* < 0.0001.

**Supplemental Figure 4.**
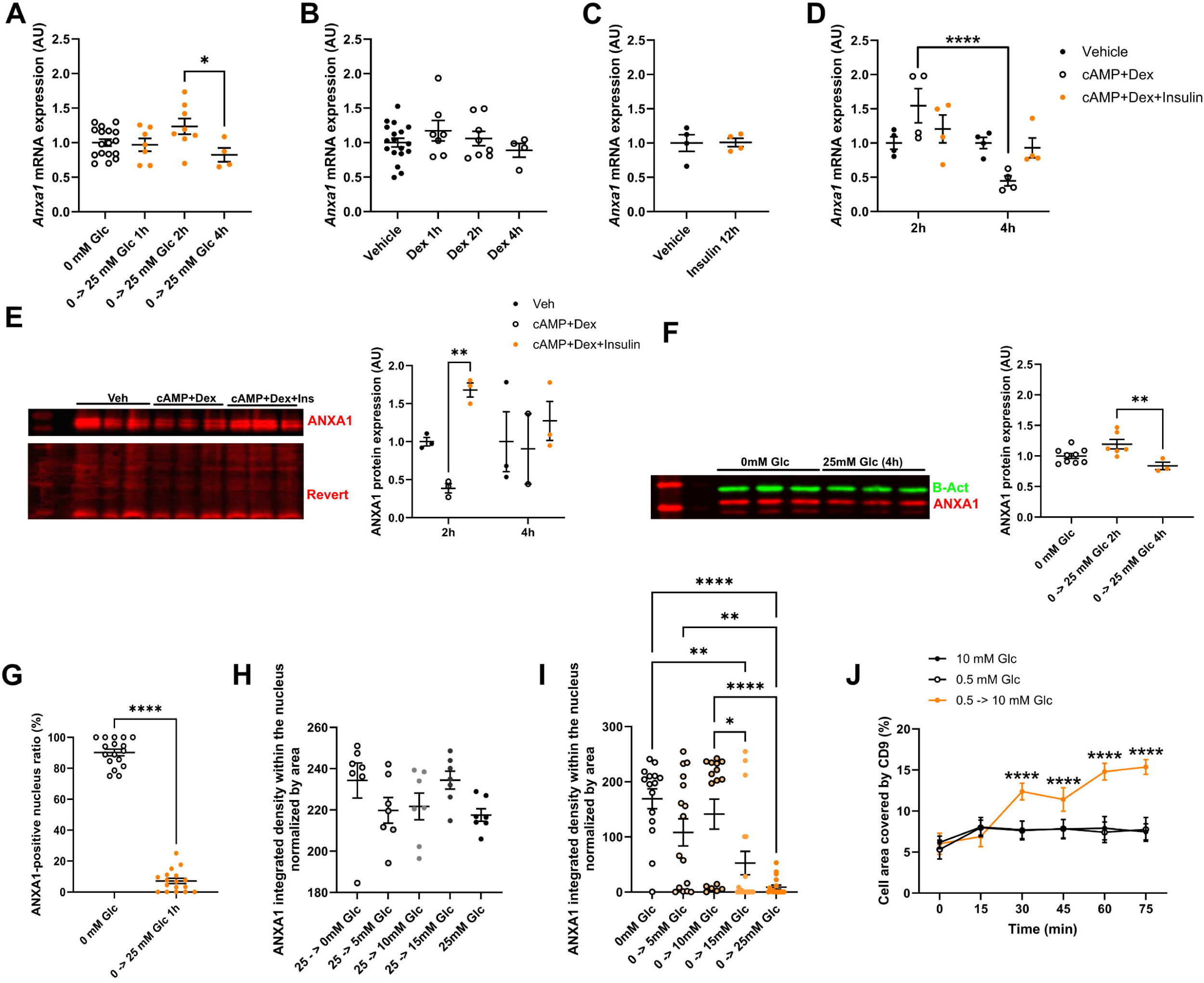
ANXA1 expression and distribution in primary tanycyte cultures. **(A-D)** *Anxa1* expression (by qPCR) in primary tanycyte cultures following glucose treatment after glucose deprivation for 24h (A) (n=4-17 per condition in 5 cultures), with dexamethasone (B) (n=4-18 per condition in 5 cultures), insulin (C) (n=4 per condition in one culture), or dexamethasone/cAMP/insulin (D) (n=4 per condition in one culture). **(E-F)** ANXA1 expression (by western blot) in primary tanycyte cultures following dexamethasone/cAMP/insulin (E) (n=3 TR per condition in one culture) or glucose treatment after glucose deprivation for 24h (F) (n=3-9 per condition in three cultures). **(G)** Percentage of tanycyte with ANXA1 staining within the nucleus following glucose treatment for 1h after glucose deprivation for 24h (n=17 pictures in one culture). **(H-I)** Glucose dose-dependent effect on ANXA1 localization while removing glucose for 1h (H) or adding glucose for 1h after glucose deprivation for 24h (I). **(J)** Time-course formation of CD9-positive vesicles following glucose treatment after glucose deprivation for 24h. 6-7 tanycytes per well were analyzed in three technical replicates in three cultures (n=9 per condition). *p < 0.05, \*\**p* < 0.01, \*\*\**p* < 0.001, \*\*\*\**p* < 0.0001.

**Supplemental Figure 5.**
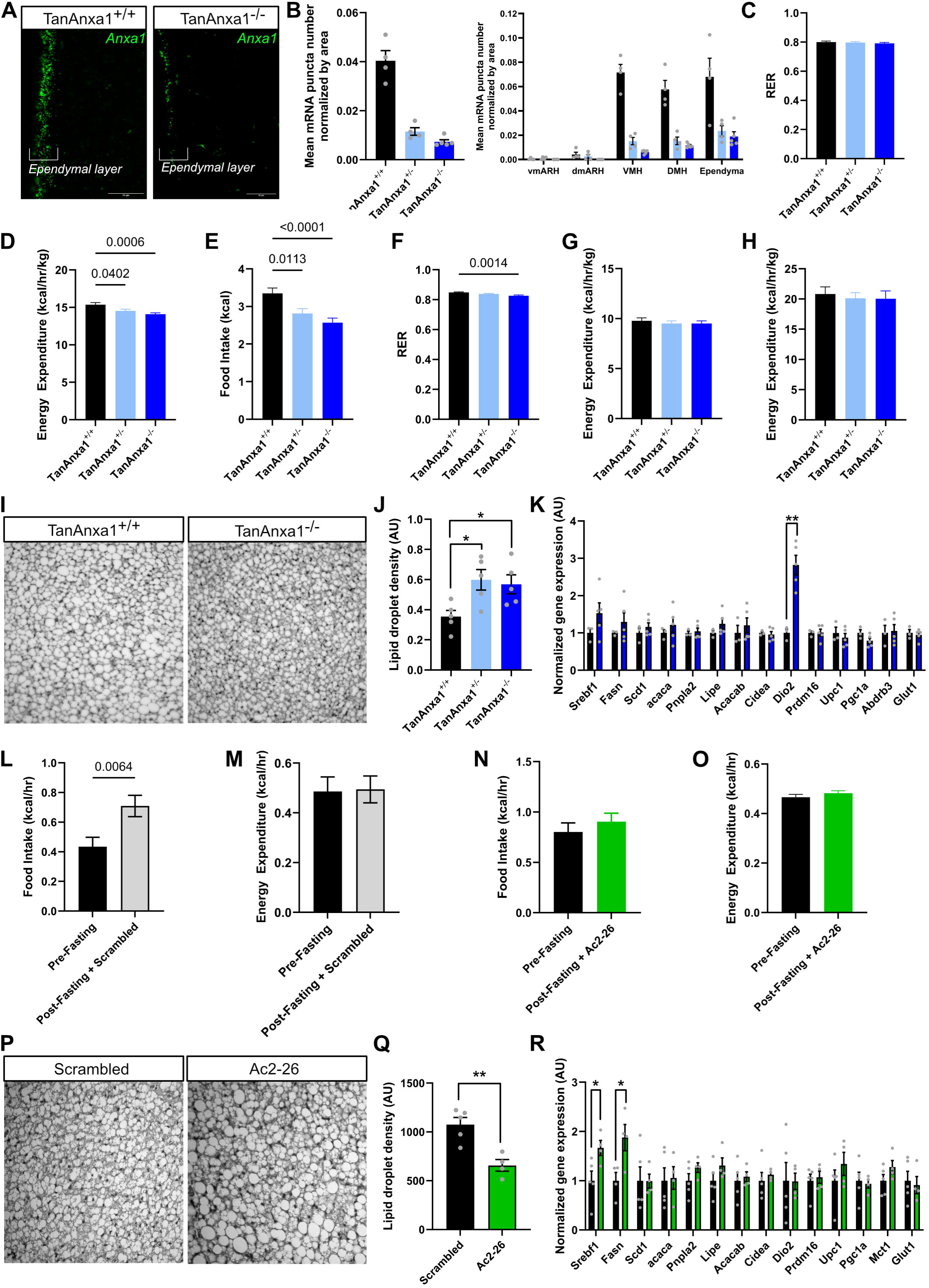
Additional physiological analysis in Ac2-26-infused or TanAnxa1^-/-^ mice. **(A-B)** Representative images (A) and quantification (B) of *Anxa1* mRNA along the ventricle in CTL *versus* TanAnxa1^−/−^ mice. **(C)** Global RER in metabolic cages in CTL *versus* TanAnxa1^−/−^ mice. **(D-F)** Energy expenditure per hour (D), cumulative food intake (E), and RER (F) in metabolic cages during refeeding after 24h fasting in CTL *versus* TanAnxa1^−/−^ mice. **(G-H)** Energy expenditure per hour at thermoneutrality (28°C, G) and 4°C (H) in CTL *versus* TanAnxa1^−/−^ mice **(I-K)** Representative images (I) and quantifications (J) of subscapular BAT histology (H&E staining) in CTL *versus* TanAnxa1^−/−^ mice; qPCR quantifications in BAT (K) in CTL *versus* TanAnxa1^−/−^ mice **(L-O)** Cumulative food intake (L, N) and energy expenditure per hour (M, O) in pre-24h-fasting *versus* post-24h-fasting in mice injected with scrambled (L-M) or Ac2-26 active peptide (N-O). **(P-R)** Representative images (P) and quantifications (Q) of subscapular BAT histology (H&E staining) in 24h-fasted mice injected with Ac2-26 active peptide; qPCR quantifications in BAT (R) in 24h-fasted mice injected with Ac2-26 active peptide. *p < 0.05, \*\**p* < 0.01, \*\*\**p* < 0.001, \*\*\*\**p* < 0.0001.

**Supplemental Figure 6.**
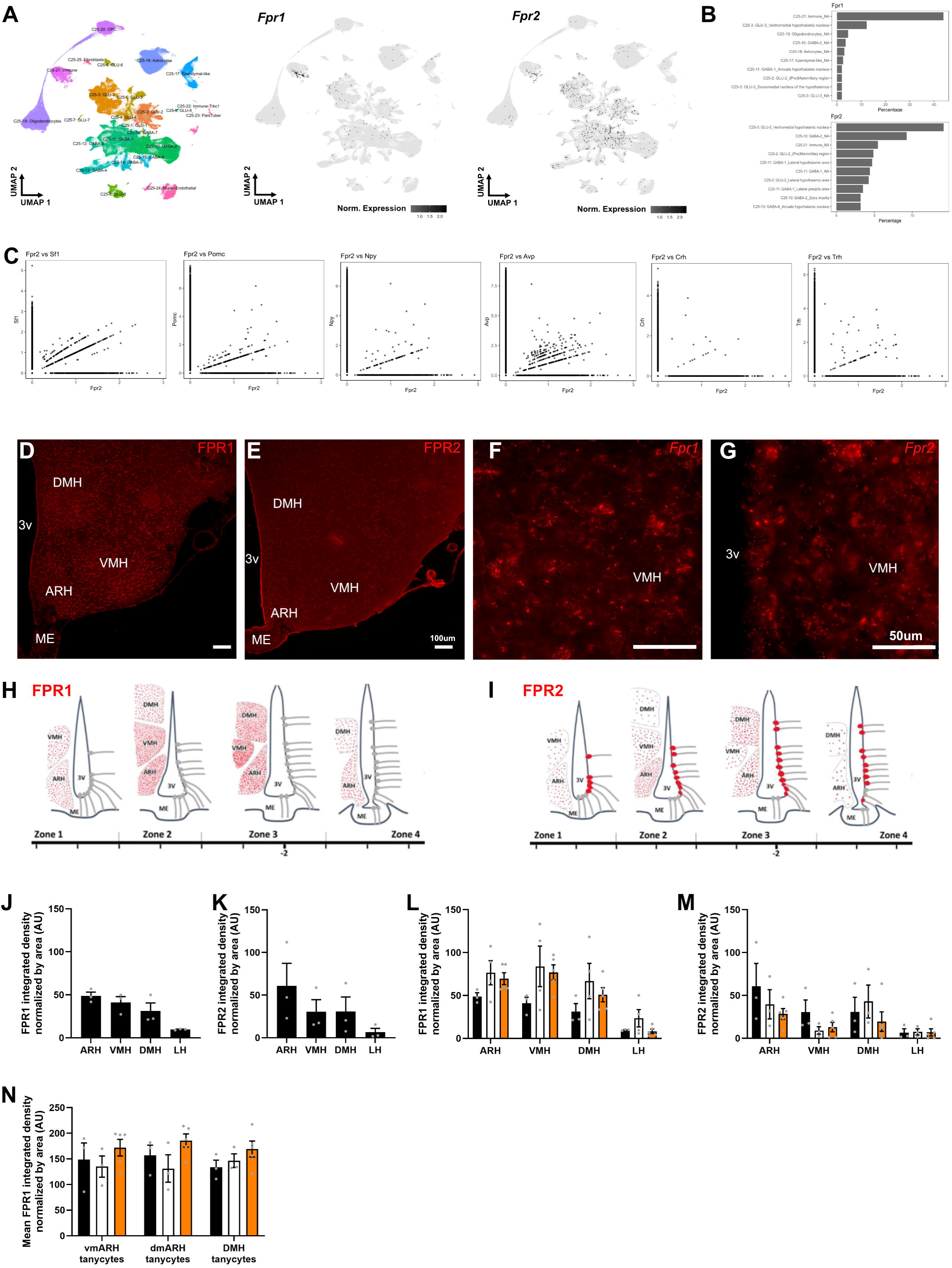
FPRs expression in the mediobasal hypothalamus. **(A)** *Fpr1* and *Fpr2* expression in the mouse hypothalamus using single-cell RNA sequencing (Mouse HypoMap). **(B)** Distribution in percentage of *Fpr1* and *Fpr2* in different subpopulations in the Mouse HypoMap^34^ **(C)** Co-expression of FPR2 with different neuronal markers (i.e., *Sf1*, *Pomc*, *Npy, Avp*, *Crh*, *Trh*) using single-cell RNA sequencing (Mouse HypoMap). **(D-G)** Representative images of FPR1 protein (D, red), FPR2 protein (E, red), *Fpr1* mRNA (F, red), and *Fpr2* mRNA (G, red) distribution in the mediobasal hypothalamus. **(H-I)** Schematic representation of FPR1 (H) and FPR2 (I) expression (red) within the mediobasal hypothalamus on the anteroposterior axis, based on *in situ* hybridization and immunohistochemistry analysis. **(J-K)** Quantitative analysis of FPR1 (J) and FPR2 (K) expression in the different nuclei (n=3 mice). **(L-M)** Quantification of FPR1 (L) and FPR2 (M) in different hypothalamic nuclei in fed, 24h-fast, and 4h-refed conditions (n=3-5 mice per condition). **(N)** Quantification of FPR2 in the tanycyte layer in fed, 24h-fast, and 4h-refed conditions (n=3-5 mice per condition). *p < 0.05, \*\**p* < 0.01, \*\*\**p* < 0.001, \*\*\*\**p* < 0.0001.

**Supplemental Figure 7.**
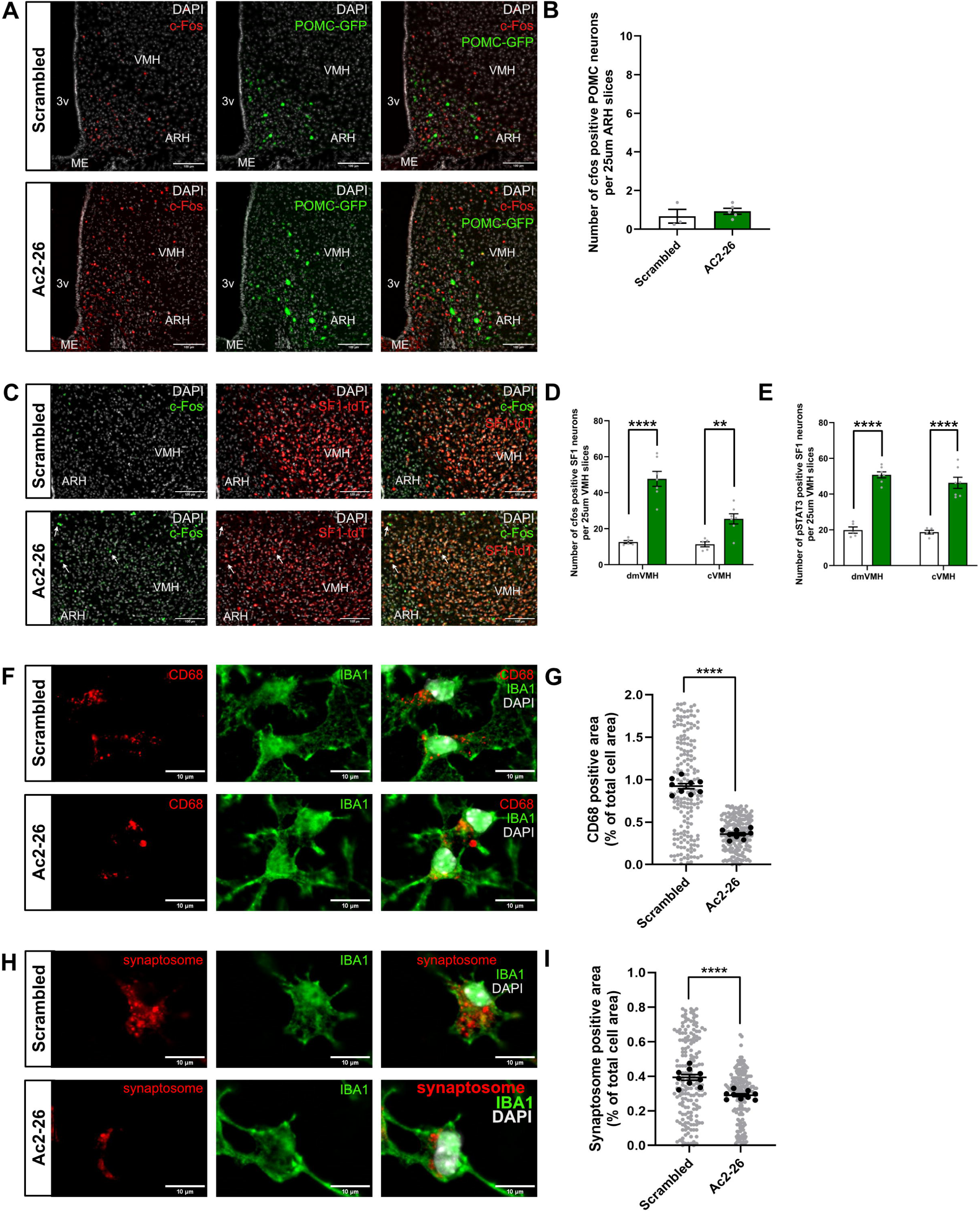
Ac2-26 infusion impact *in vivo* and *in vitro*. **(A-B)** Representative images (A) and quantifications of c-Fos activation in POMC (B) neurons following Ac2-26 i.c.v. infusion in fasted mice. **(C-E)** Representative images (C) and quantifications of c-Fos (D) and pSTAT3 (E) activation in SF1 neurons following Ac2-26 i.c.v. infusion in fasted mice. **(F-G)** Representative images (F) and quantifications (G) of CD68-positive lysosomes in primary microglia cultures following Ac2-26 treatment. **(H-I)** Representative images (H) and synaptosome assay (I) in primary microglia cultures following Ac2-26 treatment. *p < 0.05, \*\**p* < 0.01, \*\*\**p* < 0.001, \*\*\*\**p* < 0.0001.

